# LRP1 Regulates Peroxisome Biogenesis and Cholesterol Homeostasis in Oligodendrocytes and is Required in CNS Myelin Development and Repair

**DOI:** 10.1101/189563

**Authors:** Jing-Ping Lin林靚蘋, Yevgeniya A. Mironova, Peter Shrager, Roman J. Giger

## Abstract

The low-density lipoprotein related-receptor-1 (LRP1) is a large endocytic and signaling receptor. We show that *Lrp1* is required for proper CNS myelinogensis *in vivo*. Either global inducible or oligodendrocyte (OL)-lineage specific ablation of *Lrp1* impairs myelin development and adult white matter repair. In primary oligodendrocyte progenitor cells (OPCs), *Lrp1* deficiency reduces cholesterol levels and attenuates differentiation into mature OLs. Despite a strong increase in the sterol-regulatory element-binding protein-2, *Lrp1*^*-/-*^ OPCs are not able to maintain normal cholesterol levels, suggesting more global metabolic deficits. Mechanistic studies identified a decrease in peroxisomal biogenesis factor-2 and a reduction in peroxisomes localized to OL processes. Treatment of *Lrp1*^*-/-*^ OPCs with cholesterol or pharmacological activation of peroxisome proliferator-activated receptor-γ with pioglitazone is not sufficient to promote differentiation; however when combined, cholesterol and pioglitazone treatment enhance OL production. Collectively, our studies identify a novel link between LRP1, peroxisomes, and OPC differentiation during white matter development and repair.

## Introduction

In the central nervous system (CNS), the myelin-producing cell is the oligodendrocyte (OL). Mature OLs arise from oligodendrocyte progenitor cells (OPCs), a highly migratory pluripotent cell type (Rowitch and Kriegstein, 2010, Zuchero and Barres, 2013). OPCs that commit to differentiate along the OL-lineage undergo a tightly regulated process of maturation, membrane expansion, and axon myelination (Emery et al., 2009, Li and Yao, 2012, Simons and Lyons, 2013, Hernandez and Casaccia, 2015). Even after developmental myelination is completed, many OPCs persist as stable CNS resident cells that participate in normal myelin turnover and white matter repair following injury or disease (Franklin and Ffrench-Constant, 2008, Fancy et al., 2011).

LRP1 is a member of the LDL receptor family with prominent functions in endocytosis, lipid metabolism, energy homeostasis, and signal transduction (Boucher and Herz, 2011). *Lrp1* is broadly expressed in the CNS and abundantly found in OPCs (Zhang et al., 2014, Auderset et al., 2016). Global deletion of *Lrp1* is embryonically lethal (Herz et al., 1992) and conditional deletion revealed numerous tissue specific functions in neural and non-neural cell types (Lillis et al., 2008). In the PNS, *Lrp1* regulates Schwann cell survival, myelin thickness, and morphology of Remak bundles (Campana et al., 2006, Mantuano et al., 2010, Orita et al., 2013). In the CNS, *Lrp1* influences neural stem cell proliferation (Auderset et al., 2016), synaptic strength (Nakajima et al., 2013, Gan et al., 2014), axonal regeneration (Stiles et al., 2013, Yoon et al., 2013, Landowski et al., 2016), and clearance of amyloid beta (Liu et al., 2010, Zlokovic et al., 2010, Kanekiyo and Bu, 2014, Kim et al., 2014). Recent evidence shows that neurospheres deficient for *Lrp1* produce more GFAP^+^ astrocytes at the expense of O4^+^ OLs and TuJ1^+^ neurons (Hennen et al., 2013, Safina et al., 2016). Whether LRP1 is required for proper CNS myelinogenesis, nerve conduction, or repair of damaged adult CNS white matter, however, has not yet been examined. Moreover, the molecular basis of how LRP1 influences OPC differentiation remains poorly understood.

LRP1 is a large type 1 membrane protein comprised of a ligand binding 515-kDa α chain non-covalently linked to an 85-kDa β chain that contains the transmembrane domain and cytoplasmic portion. Through its α chain, LRP1 binds over 40 different ligands with diverse biological functions (Lillis et al., 2008, Fernandez-Castaneda et al., 2013). LRP1 mediates endocytotic clearance of a multitude of extracellular ligands (May et al., 2003, Tao et al., 2016) and participates in cell signaling, including activation of the Ras/MAPK and AKT pathways (Martin et al., 2008, Fuentealba et al., 2009, Muratoglu et al., 2010). The LRP1β chain can be processed by γ-secretase and translocate to the nucleus where it associates with transcription factors to regulate gene expression (May et al., 2002, Carter, 2007).

Here we combine conditional *Lrp1* gene ablation with ultrastructural and electrophysiological approaches to show that *Lrp1* is important for myelin development, nerve conduction, and adult CNS white matter repair. Gene expression analysis in *Lrp1* deficient OPCs identified a reduction in peroxisomal gene products. We show that *Lrp1* deficiency decreases production of peroxisomal proteins and disrupts cholesterol homeostasis. Mechanistic studies uncover a novel role for *Lrp1* in PPARγ mediated OPC differentiation, peroxisome biogenesis, and CNS myelination.

## Results

### *Lrp1* is required for proper CNS myelin development

In the early postnatal brain, *Lrp1* is broadly expressed (Zhang et al., 2014). In the OL-lineage, LRP1 protein is highly enriched in OPCs and absent in mature OLs (Auderset et al., 2016). To study the role of *Lrp1* in CNS myelination, we pursued a mouse genetic approach *in vivo*. To circumvent the early lethality of *Lrp1* global knockout mice (Herz et al., 1992), we generated *Lrp1*^*flox/flox*^;*CAG-creER™* mice (*Lrp1* iKO) that allow tamoxifen (TM)-inducible gene ablation under the *CMV immediate enhancer/β-actin* promoter (**Figure 1-figure supplement 1**). To assay whether LRP1 is required for proper CNS myelinogenesis, neonatal *Lrp1* control and iKO mice were subjected to TM injection at P5 and analyzed at P21 (**Figure 1a**). Western blot analysis of whole brain lysates revealed a ∼40% reduction for LRP1 in *Lrp1* iKO brain homogenate, indicating that mice are hypomorph rather than null for LRP1 (**Figure 1b**). The decrease in neural LRP1 leads to a simultaneous reduction of OL-lineage markers, of which 2’,3’-cyclic-nucleotide 3’-phosphodiesterase (CNP), proteolipid protein (PLP), and myelin basic protein (MBP) reach significance (**Figure 1c**). Ultrastructural analysis of P21 optic nerve cross-sections revealed hypomyelination in *Lrp1* iKO mice (**Figure 1d**). Only 49.7± 4.2% of axons are myelinated in *Lrp1* iKO mice, whereas 74.7± 2.4% of axons are myelinated in littermate controls (**Figure 1e**). Independent of *Lrp1* genotype, the majority of large caliber axons (>1 μm in diameter) is myelinated, while small caliber axons (<0.2 μm) are not myelinated. However, intermediate-to-small sized axons, 0.3-0.9 μm in caliber, are vulnerable to *Lrp1* deficiency and show hypomyelination (**Figure 1-figure supplement 2a**). To assess the insulating properties of myelin sheaths in the optic nerve, the ratio of the axonal diameter to the total fiber diameter (g-ratio) was calculated. The average g-ratio for myelinated axons in *Lrp1* control and iKO mice is 0.76± 0.001 and 0.81± 0.001, respectively (**Figure 1f and Figure 1-figure supplement 2b**). Axon density in the optic nerve of *Lrp1* iKO mice is similar to controls (**Figure 1-figure supplement 2c**). However, intermediate sized axons (0.4-0.8 μm) are less frequent and small axons (<0.4 μm) are more abundant in *Lrp1* iKO optic nerves (**Figure 1-figure supplement 2d**). Together, these studies show that global inducible ablation of *Lrp1* at P5 leads to a partial loss of LRP1 and CNS hypomyelination at P21.

**Figure 1-figure supplement 1:**
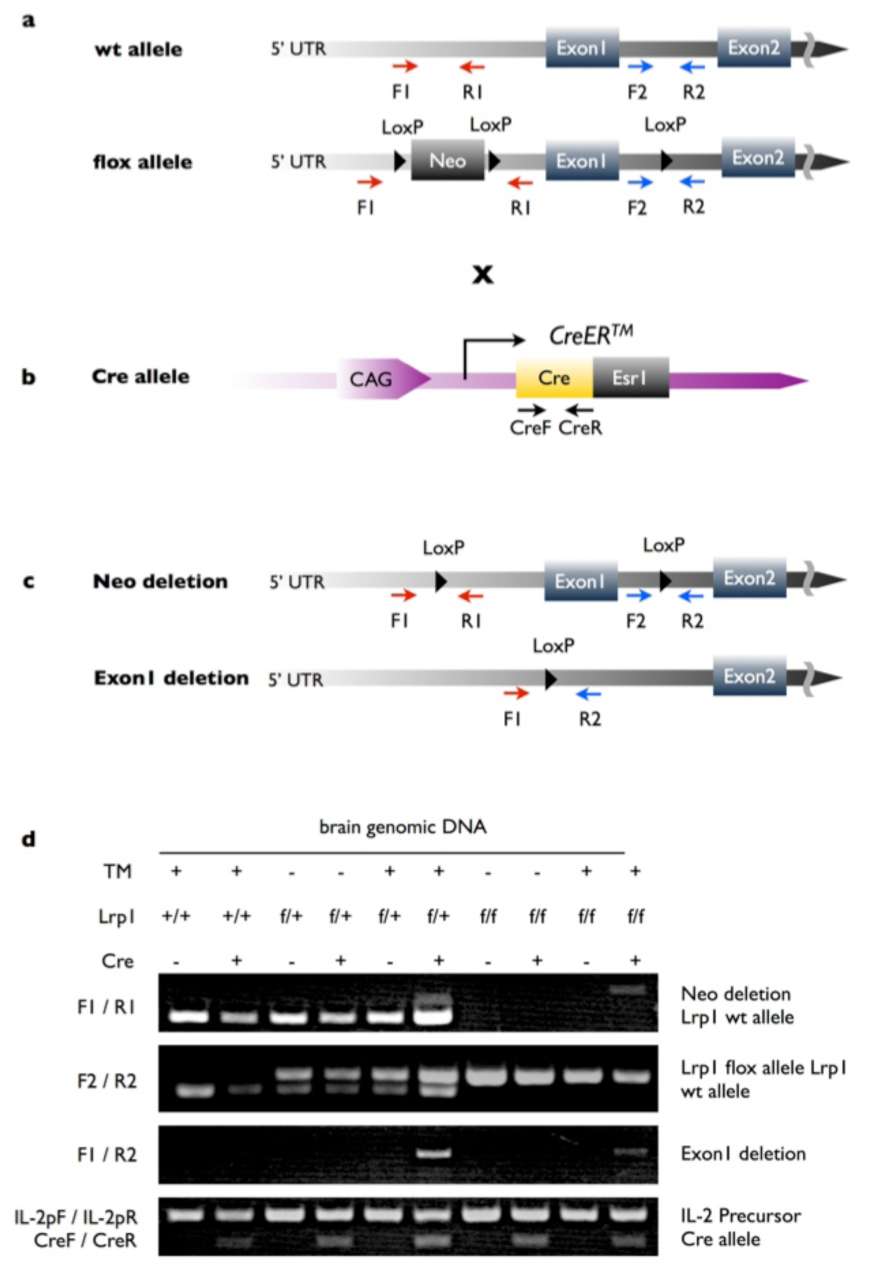
Tamoxifen induced global *Lrp1* ablation and genotyping. (a) Carton showing *Lrp1* wildtype (wt) and targeted, LoxP flanked (floxed), alleles. The location of PCR primers used for genotyping, neomycin cassette (Neo), and LoxP sites are shown. (b) For global inducible gene ablation, the *CAG-CreER™* mouse line was used, in which the Cre recombinase is fused with a TM responsive estrogen receptor (Ers1) and expressed under the control of a ubiquitous chicken β-actin-CMV hybrid (CAG) promoter. (c) Following tamoxifen (TM) administration there are two possible outcomes: deletion of the Neo cassette only or deletion of the Neo cassette and exon 1. (d) Analysis of PCR products amplified from genomic brain DNA of *Lrp1*^*+/+*^ mice with (+) or without (-) the cre allele; *Lrp1*^*flox/+*^ mice ±cre allele and ±TM treatment; *Lrp1*^*flox/flox*^ mice ±cre allele and ±TM treatment. The F1/R1 primer pair amplifies a ∼300 bp PCR product from wt *Lrp1* allele and a ∼400 bp PCR product if the Neo cassette is deleted. The F2/R2 primer pair amplifies a 291 bp PCR product from wt *Lrp1* allele and a 350 bp PCR product from *Lrp1* flox allele. The F1/R2 primer pair amplifies a ∼500 bp PCR product if exon1 in deleted. The IL-2pF/IL-2pR primer pair amplifies a 324 bp fragment and served as internal PCR quality control. The CreF/CreR primer pair amplifies a ∼200 bp PCR product from *Cre* allele.

**Figure 1:**
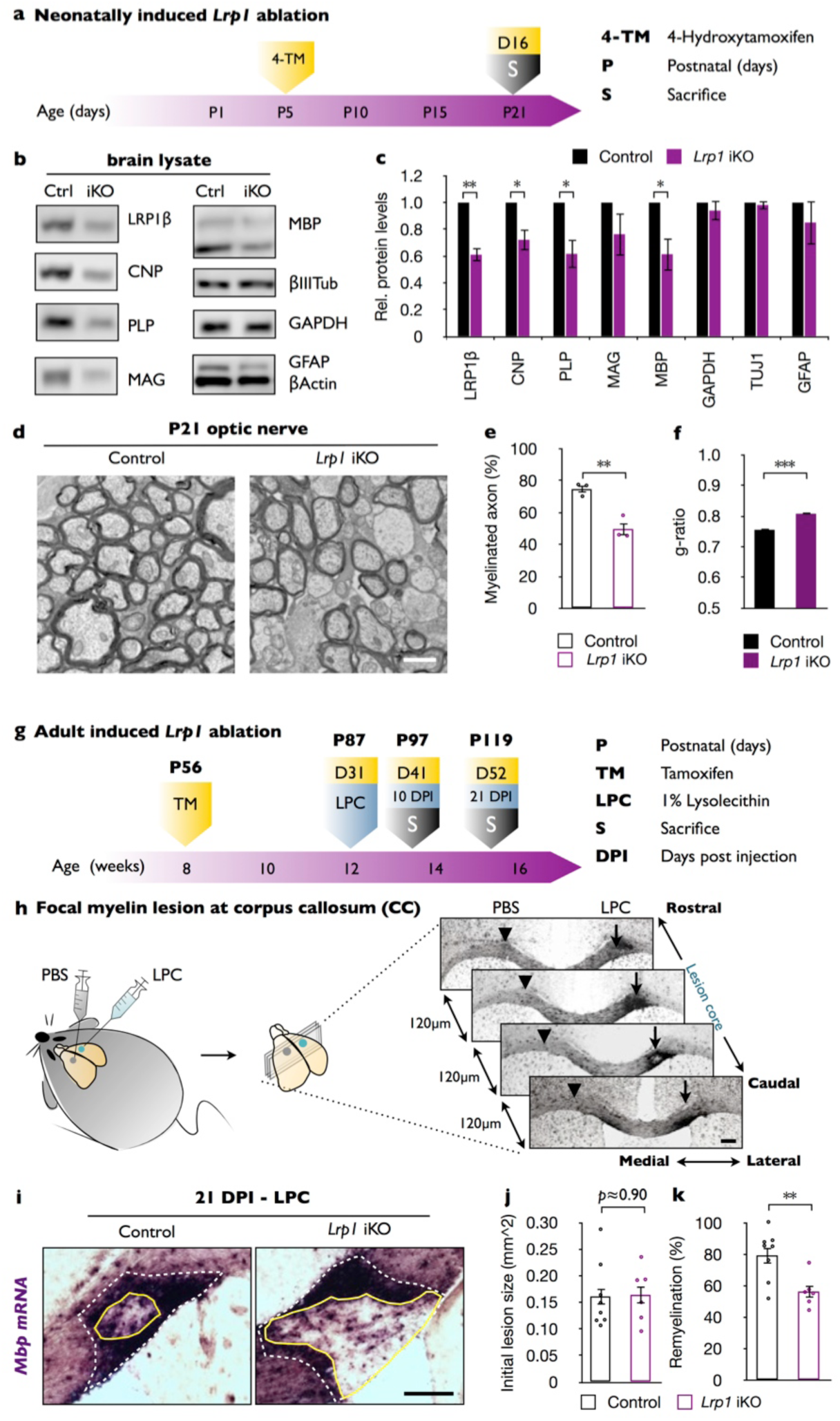
Inducible global ablation of *Lrp1* leads to CNS hypomyelination and reduced remyelination of a chemically induced white matter lesion. (a) Timeline in days showing when *Lrp1* ablation was induced and when mice were sacrificed. (b) Immunoblotting of whole brain lysates prepared from *Lrp1* control (Ctrl) and *Lrp1* inducible knockout (*Lrp1*^*flox/flox*^;*CAG-CreER*^*TM*^, *Lrp1* iKO) mice. Representative blots probed with anti-LRP1β, anti-CNP, anti-PLP, anti-MAG, anti-MBP, anti-β-III tubulin, anti-GAPDH, anti-GFAP, and anti-β-actin are shown. (c) Quantification of protein levels detected in brain lysates of *Lrp1* control (n= 3) and iKO (n= 3) mice. (d) Ultrastructural images of optic nerve cross-sections from P21 *Lrp1* control and iKO mice. Scale bar= 1μm. (e) Quantification of myelinated axons in *Lrp1* control (n= 3) and iKO (n= 3) mice. (f) Averaged g-ratio of optic nerve fibers of *Lrp1* control (n= 1932 axons, 3 mice) and iKO (n= 2461 axons, 3 mice) mice. Scale bar= 1 μm. (g) Timeline in weeks indicating when *Lrp1* ablation was induced, lysolecithin (LPC) injected, and when animals were sacrificed. (h) Cartoon showing unilateral injection of LPC in the corpus callosum (CC) and injection of PBS on the contralateral side. Coronal brain sections (series of 6, each 120 μm apart) probed for *Mbp* by *in situ* hybridization (ISH). Brain sections containing the lesion center were subjected to quantification. (i) Coronal brain sections through the CC 21 days post LPC injection (21 DPI). The outer rim of the lesion area (lesion^out^) is demarcated by the elevated *Mbp* signal (white dashed line). The non-myelinated area of the lesion is defined by the inner rim of elevated *Mbp* signal (lesion^in^) and delineated by a solid yellow line. Scale bar= 200μm. (j) Quantification of the initial lesion size (lesion^out^) in *Lrp1* control (n= 8) and iKO (n= 6) mice. (k) Quantification of the remyelinated area in *Lrp1* control (n= 8) and iKO (n=6) mice. The extent of remyelination was calculated as the percentile of (lesion^out^ - lesion^int^)/(lesion^out^). Results are shown as mean ± SEM, *p<0.05, **p<0.01, and ***p<0.001, Student’s *t*-test. For a detailed statistical report, see Figure1-source data1.

### Inducible ablation of *Lrp1* in adulthood attenuates white matter repair

To study white matter repair, 8-week-old *Lrp1* control and iKO mice were treated with i.p. TM. One month later, mice were subjected to unilateral injection of 1% lysophosphatidylcholine (LPC) into the corpus callosum and sacrificed 10 or 21 days later (**Figure 1g**). LRP1 protein levels in brain lysates of *Lrp1* iKO mice were assessed 31 and 52 days after TM injection by Western blot analysis and revealed a reduction compared to control brains (**Figure 1-figure supplement 3a**). Fluoromyelin-Green (FM-G) labeling and *in situ* hybridization (ISH) for *Mbp* transcripts of *Lrp1* control and iKO forebrain 31 days post TM injection showed comparable staining (**Figure 1-figure supplement 3b**). To examine whether *Lrp1* participates in the remyelination process following LPC-induced axon demyelination, adult mice were subjected to unilateral and focal injection of LPC into the corpus callosum. The contralateral side was injected with saline (PBS) and served as an internal control (**Figure 1h**). At 10 and 21 days post injection (DPI), mice were killed, brains extracted, and serially sectioned through the lesion area. Sections were stained with FM-G, anti-GFAP, and the nuclear dye Hoechst 33342 (**Figure 1-figure supplement 3c and 3d**). Intracranial injection of PBS led to a transient increase in GFAP, but not a reduction in FM-G staining (**Figure 1-figure supplement 3c**). Independent of *Lrp1* genotype, at 10 days following LPC injection, similar-sized white matter lesions (area devoid of FM-G labeling) and comparable astrogliosis, as assessed by GFAP staining, were noted (**Figure 1-figure supplement 3d**). At 21 DPI however, astrogliosis was reduced and the lesion area was significantly smaller in LPC injected *Lrp1* control mice compared to iKO mice (**Figure 1-figure supplement 3d**). For quantification of lesion repair, serial sections were stained with cRNA probes specific for the OPC/OL markers *Pdgfrα*, *Mag*, *Plp1*, and *Mbp*. The LPC lesion was readily detected by the upregulation of *Pdgfrα*, *Mag*, *Plp1*, and *Mbp* transcripts (**Figure 1-figure supplement 3e and 3f**). No changes for any of these transcripts were observed on the PBS injected side (**Figure 1-figure supplement 3e**). Because *Mbp* mRNA is strongly upregulated in myelin producing OLs and transported into internodes (Ainger et al., 1993), we used *Mpb in situ* hybridization on serial sections to find the center of the white matter lesion. The center was defined as the section with the largest circumference of the intensely labeled *Mbp*^*+*^ area (**Figure 1h**). The extent of initial white matter lesion, the outer rim of elevated *Mbp* labeling (white dotted line), was comparable between *Lrp1* control and iKO mice (**Figure 1l**). However, the area that failed to undergo repair, the inner rim of elevated *Mbp* labeling (yellow solid line), was larger in *Lrp1* iKO mice (**Figure 1i**). Quantification of lesion repair revealed a significant decrease in *Lrp1* iKO mice compared to controls (**Figure 1k**). As an independent assessment, serial sections through the lesion were stained for *Pdgfrα*, *Plp1,* and *Mag* transcripts and revealed fewer labeled cells within the lesion (**Figure 1-figure supplement 3g**). Together these findings indicate that in adult mice, *Lrp1* is required for the timely repair of a chemically induced white matter lesion.

**Figure 1-figure supplement 2:**
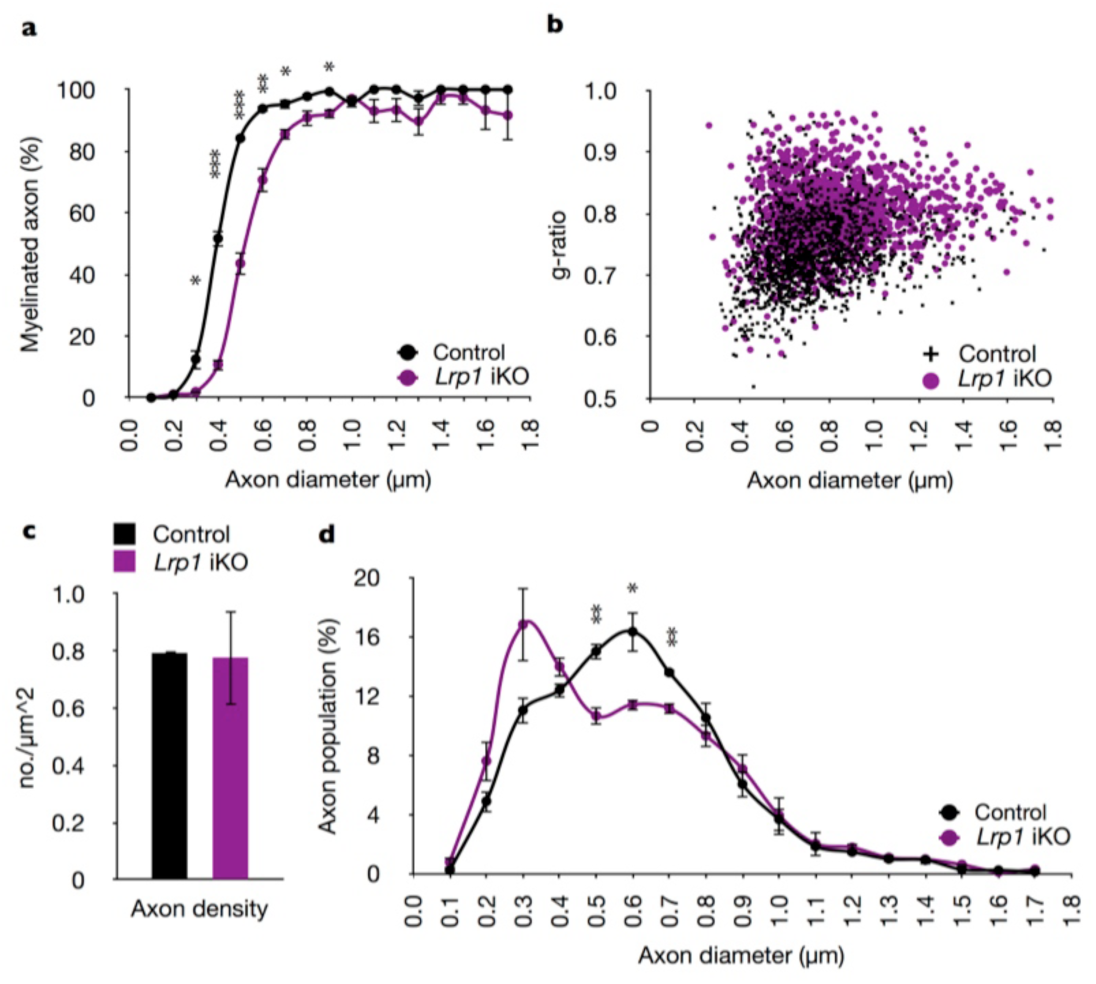
Global inducible ablation of *Lrp1* in neonatal mice leads to CNS hypomyelination and reduced axon caliber. (a) Graph showing the percentage of myelinated axons in the optic nerve of P21 mice as a function of axon caliber. Axon calibers of *Lrp1* control (n= 3) and iKO (n= 3) mice were binned into 9 groups of 0.2 μm intervals, ranging from 0.1 to 1.7 μm. The percentile of myelination is not significantly different for axons <0.3μm, significantly different for axons between 0.3-0.9μm, and not significantly different for axons between 1.0-1.7μm. (b) Scatter plot showing the distribution of g-ratios for individual fibers in the optic nerve of P21 *Lrp1* control (n= 2461 axons, 3 mice) and iKO (n= 1934 axons, 3 mice). (c) Quantification of axon density per μm^2^ in P21 optic nerve cross sections of *Lrp1* control (n= 3) and iKO (n= 3) mice. (d) Morphometric assessment of axon caliber distribution in P21 optic nerves of *Lrp1* control (n= 3) and iKO (n= 3) mice. Axon diameters were measured on electron microscopy images and quantification revealed a shift toward smaller-sized axons in *Lrp1* iKO mice. Results are presented as the mean ± SEM, \**p*<0.05, \*\**p*<0.01, and \*\*\**p*<0.001, Student’s *t*-test. For a detailed statistical report, see Figure1-source data1.

**Figure 1-figure supplement 3:**
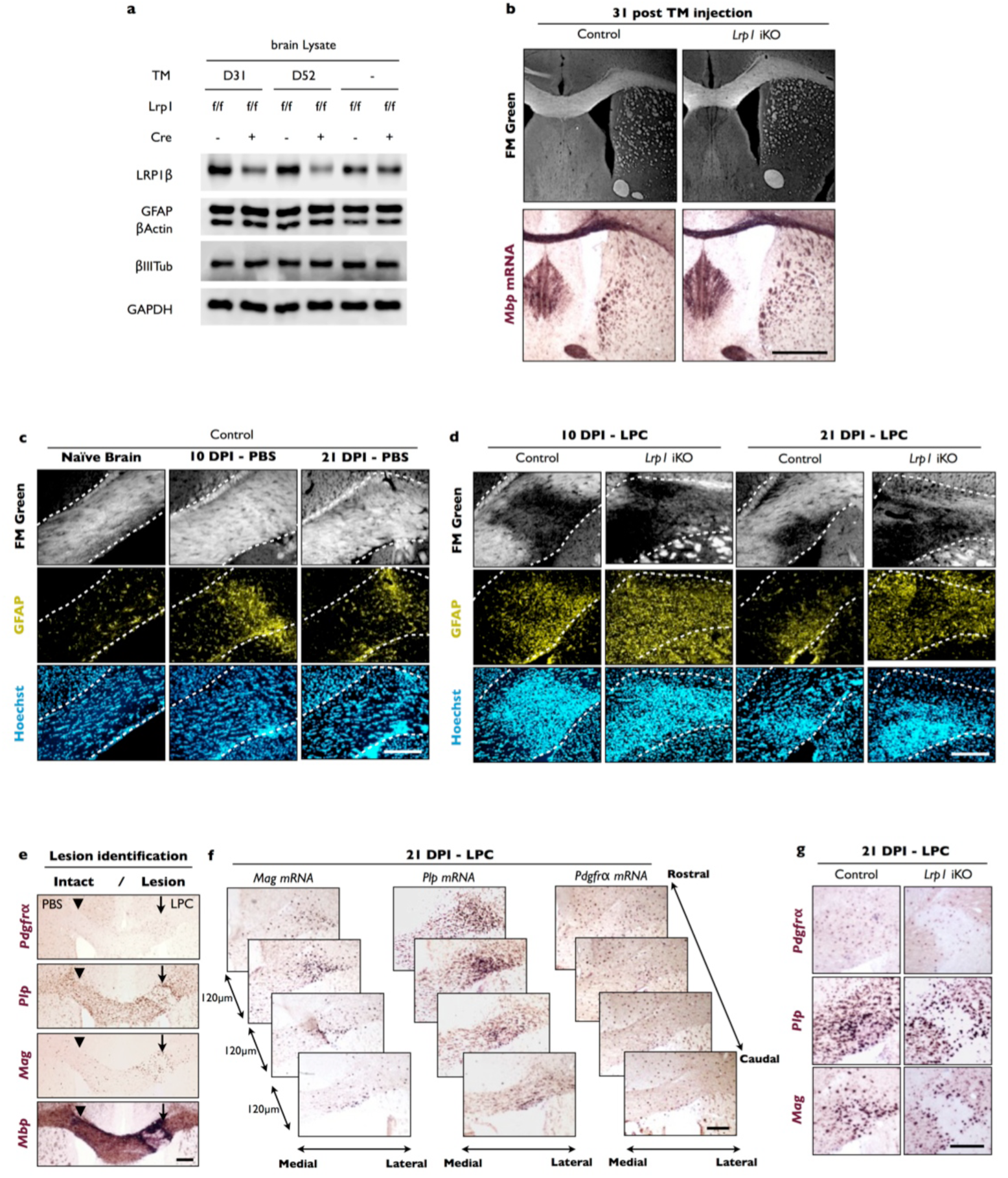
*Lrp1* is important for white matter repair. (a) Immunoblots of whole brain lysates prepared from *Lrp1*^*flox/flox*^; *CAG-creER™* mice 31 and 52 days after TM (+) or vehicle (-) treatment. Representative blots probed with anti-LRP1β, anti-GFAP, anti-β-actin, anti-β-III tubulin, and anti-GAPDH. (b) Coronal-sections of adult *Lrp1* control and iKO mice 31 days after i.p. TM administration. Sections were stained with FM Green or probed for *Mpb* mRNA by *in situ* hybridization. Scale bar= 1mm. (c) Coronal forebrain sections of *Lrp1* control mice without injection (naïve) or PBS injection in the corpus callosum. At 10 days post injection (10 DPI) and 21 DPI of PBS brains were sectioned and stained with FM Green, anti-GFAP and Hoechst dye33342. The white dotted lines demarcate the corpus callosum. (d) Coronal forebrain sections of *Lrp1* control and iKO mice at 10 DPI and 21 DPI of LPC stained with FM Green, anti-GFAP, and Hoechst dye33342. White dotted lines demarcate the corpus callosum. Scale bar= 200μm. (e) Serial coronal-sections of adult brain after PBS and LPC injection in the corpus callosum were probed for *Pdgfrα*, *Plp1*, *Mag*, and *Mbp* mRNA to identify the lesion area and to examine gene expression changes in the OL lineage. The PBS injection site is marked by an arrowhead and the LPC injection site is marked by an arrow. Scale bar= 200μm. (f) Serial brain sections of adult *Lrp1* control mice injected with LPC at 21 DPI. Sections that contain lesion area (120 μm apart) were probed for *Pdgfrα*, *Plp1*, and *Mag* mRNA expression. Scale bar= 200μm. (g) Coronal sections through the corpus callosum of *Lrp1* control and *Lrp1* iKO mice injected with LPC. Serial sections were probed for *Pdgfrα*, *Plp1*, and *Mag* mRNA expression. Scale bar= 200μm.

### Ablation of *Lrp1* in the OL-lineage causes hypomyelination

To demonstrate an OL-lineage specific function for *Lrp1* during CNS myelination, we generated *Lrp1*^*flox/flox*^;*Olig2-cre* mice (*Lrp1* cKO^OL^) (**Figure 2-figure supplement 1a**). *Lrp1* cKO^OL^ pups are born at the expected Mendelian frequency and show no obvious abnormalities at the gross anatomical level. LRP1 protein levels in the brains of P10, P21, and P56 *Lrp1* control and cKO^OL^ mice were analyzed by Western blot analysis and revealed a partial loss of LRP1β (**Figure 2-figure supplement 1b**). The partial loss of LRP1β in brain lysates of OL-lineage specific *Lrp1* cKO^OL^ mice is due to *Lrp1* expression in several other neural cell types (Zhang et al., 2014).

To examine whether *Lrp1* cKO^OL^ mice exhibit defects in myelination, optic nerves were isolated at P10, the onset of myelination; at P21, near completion of myelination; and at P56, when myelination is thought to be largely completed. Ultrastructural analysis at P10 revealed no significant difference in myelinated axons between *Lrp1* control (17± 6%) and cKO^OL^ (7± 2%) optic nerves (**Figure 2a and 2b**). At P21 and P56, the percentile of myelinated axons in the optic nerve of cKO^OL^ mice (49± 4% and 66± 5%, respectively) is significantly reduced compared to controls (70± 2% and 88± 1%, respectively) (**Figure 2a and 2b**). Similarly to *Lrp1* iKO mice (**Figure 1-figure supplement 2a**), in *Lrp1* cKO^OL^ mice intermediate to small sized axons, 0.3-0.9μm in caliber, are more vulnerable to hypomyelination (**Figure 2-figure supplement 1c, 1f, and 1i**). As an independent assessment of fiber structure, the g-ratio was determined. At P10, P21, and P56 the average g-ratio of *Lrp1* cKO^OL^ optic fibers is significantly larger than in age-matched *Lrp1* control mice (**Figure 2c and Figure 2-figure supplement 1d, 1g, and 1j**). While axon density in the optic nerve of *Lrp1* cKO^OL^ mice is similar to littermate controls at all three time points examined, there is a shift in axon caliber, similar to *Lrp1* iKO mice (**Figure 2-figure supplement 1e, 1h, 1k, and Figure 1-figure supplement 2d**). Comparable to P21 *Lrp1* iKO mice (**Figure 1b and 1c**), Western blot analysis of adult *Lrp1* cKO^OL^ brain lysates revealed a significant reduction in CNP, MAG, and MBP (**Figure 2-figure supplement 1l and 1m**). Together, these studies show that in the OL lineage *Lrp1* functions in a cell-autonomous manner and is required for proper CNS myelinogenesis.

**Figure 2:**
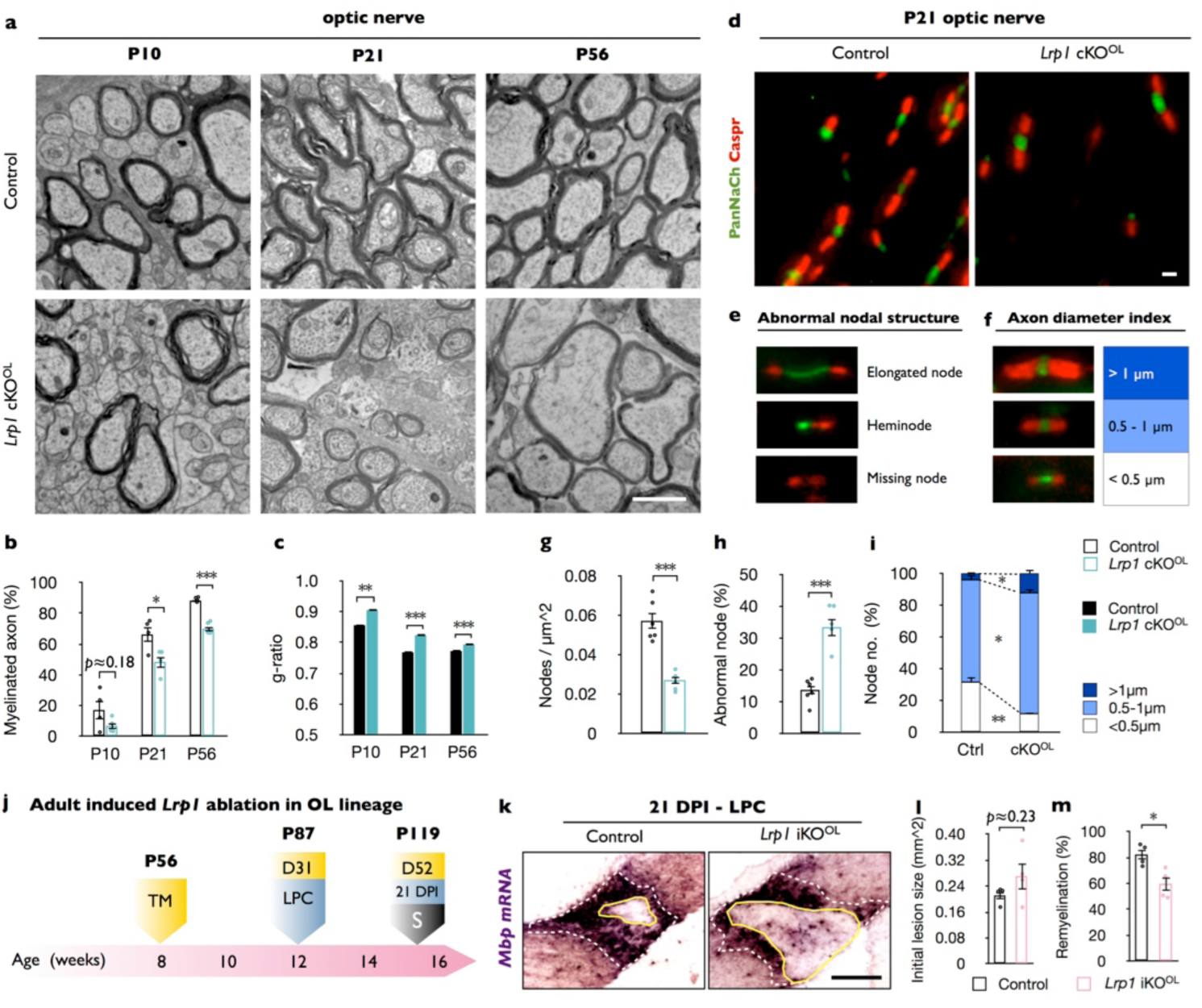
*Lrp1* ablation in the oligodendrocyte (OL)-lineage leads to hypomyelination, nodal defects, and reduced myelin repair. (a) Ultrastructural images of optic nerve cross-sections from P10, P21, and P56 control and *Lrp1*^*flox/flox*^;*Olig2-Cre* conditional knockout mice (*Lrp1* cKO^OL^). Scale bar= 1μm. (b) Quantification of myelinated axons in the optic nerve of *Lrp1* control (n= 4 mice per time point) and cKO^OL^ (n= 4 mice per time point) mice. (c) Averaged g-ratio of *Lrp1* control and cKO^OL^ optic nerve fibers from 4 mice per time points in each group. At P10, n= 488 axons for control and n= 261 axons for cKO^OL^; at P21, n= 1015 axons for control and n= 997 axons for cKO^OL^; at P56, n= 1481 axons for control and n= 1020 axons for cKO^OL^ were quantified. (d) Nodes of Ranvier in P21 optic nerves of *Lrp1* control and cKO^OL^ mice were labeled by anti-PanNaCh (green, node) and anti-Caspr (red, paranode) staining. Scale bar= 1μm. (e) Nodal defects detected include elongated node, heminode, and missing node (Na^+^ channels absent). (f) Representative node staining categorized by axon diameter. (g) Quantification of nodal density in *Lrp1* control (n= 6) and cKO^OL^ (n= 5) optic nerves. (h) Quantification of abnormal nodes of Ranvier in *Lrp1* control (n= 6) and cKO^OL^ (n= 5) optic nerves. (i) Quantification of nodal frequency in axons with large, intermediate, and small caliber for *Lrp1* control (n= 6) and cKO^OL^ (n= 5) optic nerves. (j) Timeline in weeks showing when OL-lineage specific *Lrp1* ablation (*Lrp1*^*flox/flox*^;*Pdgfrα-CreER*^*TM*^, *Lrp1* iKO^OL^) was induced, LPC injected, and when animals were sacrificed. (k) Coronal brain sections through the CC at 21 days post LPC injection of *Lrp1* control and iKO^OL^ mice. The initial lesion area is demarcated by a white dashed line. A solid yellow line delineates the non-myelinated area. Scale bar= 200μm. (l) Quantification showing the initial lesion size in *Lrp1* control (n= 4) and iKO^OL^ (n= 4) mice. (m) Quantification of the remyelinated area in *Lrp1* control (n= 4) and iKO^OL^ (n= 4) mice. Results are shown as mean ± SEM, *p<0.05, **p<0.01, and ***p<0.001, Student’s *t*-test. For a detailed statistical report, see Figure2-source data1.

**Figure 2-figure supplement 1:**
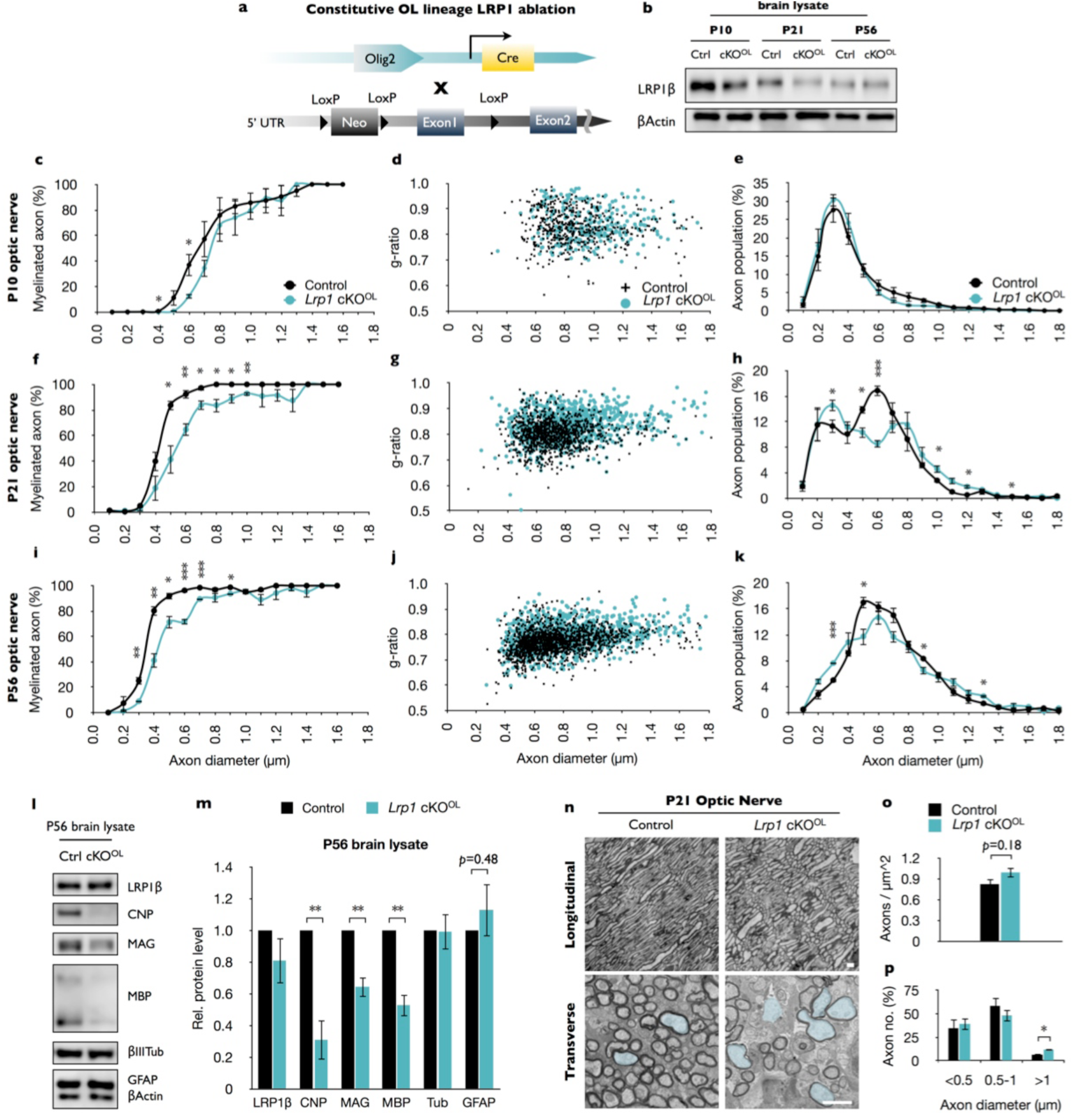
*Lrp1* ablation in the OL lineage leads to CNS hypomyelination. (a) *Lrp1*^*flox/flox*^ mice were crossed with *Olig2-cre* mice to conditionally ablate *Lrp1* in OL lineage (cKO^OL^). (b) Immunoblotting of whole brain lysates prepared from P10, P21, and P56 of *Lrp1* control(Ctrl) and cKO^OL^ mice. Representative blots probed with anti-LRP1β and anti-β-Actin. (c, f, and i) Graphs show the percentage of myelinated axons P10, P21 and P56 as a function of axon caliber for *Lrp1* control (n= 4 for each time point) and cKO^OL^ (n= 4 for each time point) mice. Axon calibers were binned into 9 groups of 0.2 μm intervals, ranging from 0.1 to 1.7 μm. (d, g, and j) Scatter plot showing the distribution of g-ratios for individual fibers in the optic nerve at P10, P21 and P56 of *Lrp1* control and cKO^OL^ mice. P10, n= 488 axons from *Lrp1* control mice and n= 261 axons from cKO^OL^ mice; P21, n= 1015 axons from *Lrp1* control mice and n= 997 axons from 4 cKO^OL^ mice; P56, n= 1481 axons from *Lrp1* control mice and n= 1020 axons from 4 cKO^OL^ mice. (e, h, and k) Morphometric assessment of axon caliber distribution in P10, P21 and P56 optic nerves of *Lrp1* control (n= 4) and cKO^OL^ (n= 4) mice. Measurements of axon diameter were made from electron microscopy images. In *Lrp1* cKO^OL^ optic nerves, the population at smaller-sized (0.2-0.4μm) axons is reduced at P21 and P56. In addition, there is shift towards larger sized axons at P21 in *Lrp1* cKO^OL^ optic nerves (h). (l) Immunoblotting of whole brain lysates prepared from *Lrp1* control (Ctrl) and cKO^OL^ mice. Representative blots probed with anti-LRP1β, anti-CNP, anti-MAG, anti-MBP, anti-β-III tubulin, anti-GFAP, and anti-β-actin. (m) Quantification of protein levels detected by Western blotting of *Lrp1* control (n= 3) and cKO^OL^ (n= 3) brain lysates. (n) Electron microscopy images of optic nerve cross- and longitudinal-sections acquired from P21 *Lrp1* control and cKO^OL^ mice. Axons that are >1μm in diameter are colored in light blue. Scale bar= 1μm. (o) Quantification of axon density in the P21 optic nerve for *Lrp1* control (n= 4) and cKO^OL^ (n=4) mice. (p) Quantification of axons in the optic nerve that are smaller than < 0.5μm, between 0.5-1.0μm and larger than 1μm for *Lrp1* control (n= 4) and cKO^OL^ (n= 4) mice. Results are presented as the mean ± SEM, \**p*<0.05, \*\**p*<0.01, and \*\*\**p*<0.001, Student’s *t*-test. For a detailed statistical report, see Figure2-source data1.

### Loss of *Lrp1* in the OL-lineage causes defects in nodes of Ranvier and faulty nerve conduction

To examine whether *Lrp1* in OLs is required for nodal or paranodal organization, optic nerve sections of P21 *Lrp1* control and cKO^OL^ mice were immunostained for sodium channels (PanNaCh) and the paranodal axonal protein (Caspr). Nodal density, the number of PanNaCh^+^ clusters in longitudinal optic nerve sections is significantly reduced in *Lrp1* cKO^OL^ mice (**Figure 2d and 2g**). In addition, an increase in nodal structural defects, including elongated nodes, heminodes, and nodes in which sodium channel staining is missing, was observed in mutant nerves (**Figure 2e**). Quantification of nodal structural defects revealed an increase from 13.7± 1.3% in *Lrp1* control mice to 33.4± 2.9% in cKO^OL^ optic nerves (**Figure 2h**). Of the total number of PanNaCh^+^ nodes quantified (including elongated nodes and heminodes), a greater fraction is associated with large (>1 μm) and intermediate (0.5-1 μm) caliber axons in *Lrp1* cKO^OL^ mice, while the number of nodes associated with small (<0.5 μm) caliber axons is significantly reduced in mutants (**Figure 2i**). The total number of optic nerve axons does not change between *Lrp1* control and cKO^OL^ mice, but the fraction of large diameter axons is significantly increased in mutants (**Figure 2-figure supplement 1n-1p**). To assess whether structural defects observed in optic nerve myelin, nodes of Ranvier, and axon caliber distribution in *Lrp1* cKO^OL^ mice are associated with impaired nerve conduction, we used electrophysiological methods to measure compound action potentials (CAPs) in acutely isolated nerves (**Figure 2-figure supplement 2a**). Analysis of CAP recordings identified 4 peaks with distinct latencies in P21 optic nerves. Representative traces of *Lrp1* control and cKO^OL^ nerve recordings were fitted as the sum of four Gaussians (**Figure 2-figure supplement 2b**). Peaks 1, 2, and 3 are likely obtained from myelinated axons and Peak 4 represents a population of slow conducting, possibly non-myelinated axons known to be present in the P21 optic nerve(Mironova et al., 2016). A shift in conduction to the right, which reflects an increase in the relative contribution of unmyelinated axons to the CAP and a reduction in amplitudes were observed in *Lrp1* cKO^OL^ nerves (**Figure 2-figure supplement 2c-2f**). Changes in electrophysiological properties in *Lrp1* cKO^OL^ nerves fit well with defects at the ultrastructural level and aberrant node assembly. Taken together, OL-lineage specific ablation of *Lrp1* leads to impaired nerve conduction.

**Figure 2-figure supplement 2:**
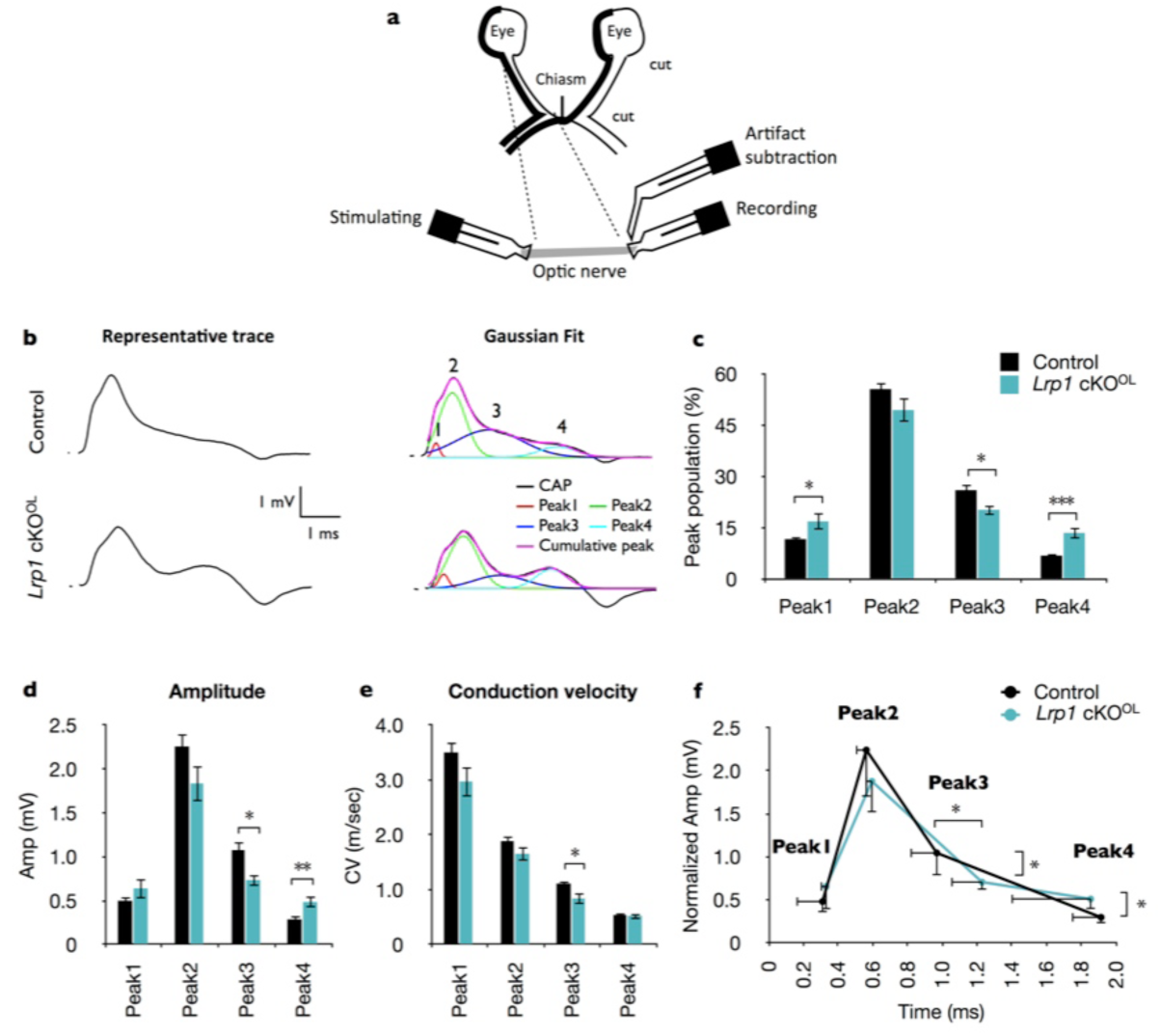
Loss of *Lrp1* in the OL lineage leads to faulty nerve conduction. (a) Scheme depicting the orientation of an optic nerve prepared for compound action potential (CAP) recording. Positions of the stimulating electrode, the recording electrode, and artifact subtraction electrode are shown. (b) Left: representative raw CAP traces of P21 optic nerves. Right: For each recording, traces were fitted with 4 Gaussians representing peak 1 (red), peak 2 (green), peak 3 (blue), peak 4 (cyan), and the sum of the four peaks (magenta). (c) The distribution of peak populations in *Lrp1* control and cKO^OL^ mice. (d) Quantification of amplitudes (mV) of peaks 1, 2, 3 and 4 in *Lrp1* control and cKO^OL^ optic nerves. (e) Quantification of conduction velocities (m/sec) of peaks 1, 2, 3 and 4 in *Lrp1* control and cKO^OL^ optic nerves. (f) Reconstituted averaged peak1-4 amplitude as a function of time. For *Lrp1* control mice n= 21 nerves from 14 mice and for cKO^OL^ n= 9 nerves from 7 mice. Results are presented as the mean ±SEM, \**p*<0.05, \*\**p*<0.01, and \*\*\**p*<0.001, Student’s *t*-test. For a detailed statistical report, see Figure2-source data1.

### OL-lineage specific ablation of *Lrp1* impairs timely repair of damaged white matter

To determine the cell autonomy of *Lrp1* in adult white matter repair, we generated mice that allow inducible *Lrp1* ablation selectively in OPCs in adult mice. Inducible gene ablation was necessary to rule out potential confounding effects on repair, originating from developmental white matter defects observed in adult *Lrp1* cKO^OL^ mice (**Figure 2a**). For repair studies, *Lrp1*^*flox/flox*^;*PDGFRα-creER™* (*Lrp1* iKO^OL^) mice were generated and gene ablation was induced by TM administration at 8 weeks of age (**Figure 2j**). One month later, mice were subjected to stereotaxic injection of LPC into the corpus callosum. The contralateral side was injected with PBS and served as a negative control. Twenty-one days post injection (21 DPI) brains were collected and analyzed. Detection of the initial white matter lesion and quantification of the extent of axon re-myelination was carried out as shown above (**Figure 1h and 1i**). The initial size of the LPC inflicted white matter lesion is comparable between *Lrp1* control and iKO^OL^ mice (**Figure 2k and 2l**). The extent of lesion repair was significantly decreased in *Lrp1* iKO^OL^ mice (**Figure 2m**), demonstrating a cellautonomous role for *Lrp1* in the OL-lineage for the timely repair of a white matter lesion.

**Figure 3:**
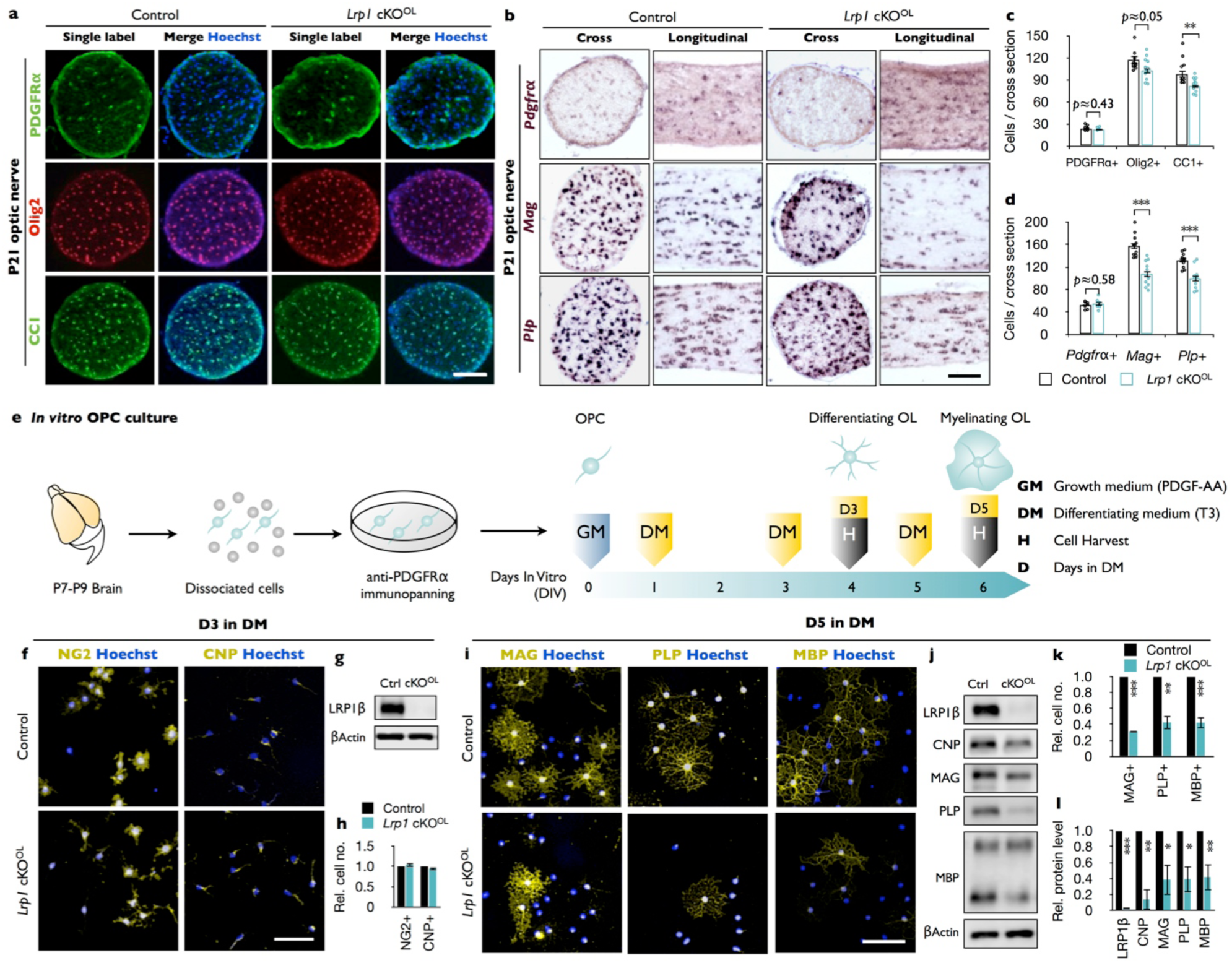
Loss of *Lrp1* in the OL-lineage attenuates OL differentiation. (a) Cross-sections of *Lrp1* control and cKO^OL^ mice stained with anti-PDGFRα (OPC marker), anti-Olig2 (pan-OL marker), anti-CC1 (mature OL marker), and Hoechst dye33342. Scale bar= 100μm. (b) Cross- and -longitudinal sections of *Lrp1* control and cKO^OL^ optic nerves probed for *Pdgfrα*, *Mag*, and *Plp* mRNA expression. Scale bar= 100μm. (c) Quantification of labeled cells per nerve cross-section. Anti-PDGFRα, n= 8 for controls and n= 6 for cKO^OL^ mice; anti-Olig2 and anti-CC1, n=11 for control and n=12 for cKO^OL^ mice. (d) Quantification of labeled cells per nerve cross-section. *Pdgfrα*, n= 8 for control and n= 6 for cKO^OL^ mice; *Mag*, n= 11 for controls and n= 11 for cKO^OL^ mice; *Plp*, n= 11 for controls and n= 10 for cKO^OL^ mice. (e) Workflow for OPC isolation with timeline when growth medium (GM) or differentiation medium (DM) was added and cells were harvested. (f) OPC/OL culture after 3 days in DM stained with anti-NG2 (premyelinating marker), anti-CNP (differentiating OL marker), and Hoechst dye33342. Scale bar= 100μm. (g) Immunoblot of OL lysates prepared from *Lrp1* control and cKO^OL^ cultures after 3 days in DM probed with anti-LRP1β and anti-β-actin. (h) Quantification of NG2^+^ (n= 3 per condition) and CNP^+^ (n= 3 per condition) cells in *Lrp1* control and cKO^OL^ cultures. (i) OL cultures after 5 days in DM stained with anti-MAG, anti-PLP, and anti-MBP. Scale bar= 100μm. (j) Immunoblotting of OL lysates prepared from *Lrp1* control and cKO^OL^ cultures after 5 days in DM probed with anti-LRP1β, anti-CNP, anti-MAG, anti-PLP, anti-MBP, and anti-β-actin. (k) Quantification of MAG^+^ (n= 3 per condition), PLP^+^ (n= 3 per condition), and MBP^+^ (n= 5 per condition) cells in *Lrp1* control and cKO^OL^ cultures. (l) Quantification of protein levels in OL lysates detected by immunoblotting. Anti-LRP1, CNP, and PLP, n= 3 per condition; anti-MAG, n= 4 per condition; anti-MBP n= 5 per condition. Results are shown as mean values ±SEM, *p<0.05, **p<0.01, and ***p<0.001, Student’s *t*-test. For a detailed statistical report, see Figure3-source data1.

### Conditional ablation of *Lrp1* in the OL-lineage attenuates OPC differentiation

CNS hypomyelination in *Lrp1* cKO^OL^ mice may be the result of reduced OPC production or impaired OPC differentiation into myelin producing OLs. To distinguish between these two possibilities, optic nerve cross-sections were stained with anti-PDGFRα, a marker for OPCs, anti-Olig2 to account for all OL lineage cells in the culture, and anti-CC1, a marker for mature OLs. No change in OPC density was observed, but the number of mature OLs was significantly reduced in *Lrp1* cKO^OL^ mice (**Figure 3a and 3c**). Optic nerve ISH for *Pdgfrα*^*+*^ revealed no reduction in labeled cells in *Lrp1* cKO^OL^ mice, a finding consistent with anti-PDGFRα immunostaining. The density of *Plp* and *Mag* expressing cells, however, is significantly reduced in the optic nerve cross-sections and longitudinal-sections of *Lrp1* cKO^OL^ mice (**Figure 3b and 3d**). These studies reveal that OPCs are present at normal density and tissue distribution in *Lrp1* cKO^OL^ mice, but apparently fail to generate sufficient numbers of mature, myelin-producing OLs.

**Figure 3-figure supplement 1:**
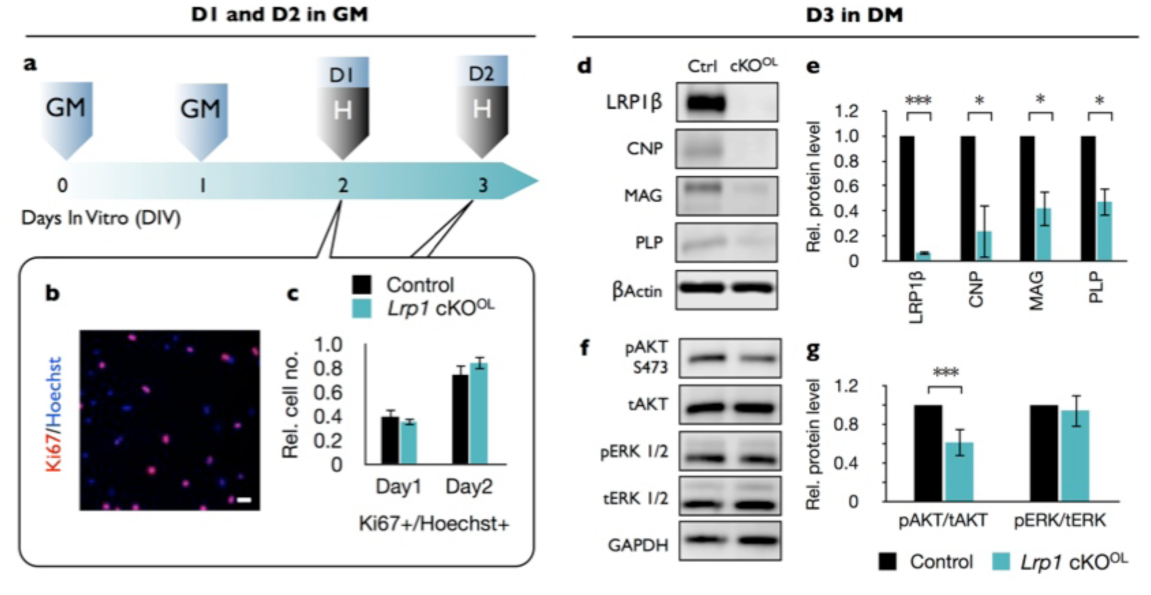
Loss of *Lrp1* does not alter OPC proliferation. (a) Timeline in days indicating when growth medium (GM) was added to cells and when cells were harvested (H) to assess proliferation. (b) OPC/OL culture after 1 or 2 days in GM stained with anti-Ki67 (proliferation marker) and Hoechst dye33342. Scale bar= 100μm. (c) Quantification of cell proliferation in OPC/OL cultures prepared from *Lrp1* control and cKO^OL^ mice. The percentile of Ki67^+^/Hoechst^+^ cells was calculated on day 1 for *Lrp1* control (n= 5) and cKO^OL^ (n= 5) cultures and on day 2 for *Lrp1* control (n= 4) and cKO^OL^ (n = 4) cultures. (d) Immunoblotting of cell lysates prepared from *Lrp1* control and cKO^OL^ OPC/OL cultures after 3 days in differentiation medium (DM). Blots were probed with anti-LRP1β, anti-CNP, anti-MAG, anti-PLP, and anti-β-actin. (e) Quantification of protein levels detected by immunoblotting per OPC/OL culture from *Lrp1* control and cKO^OL^ mice. Anti-LRP1 and anti-MAG, n=3 for per condition; anti-CNP and anti-PLP, n=4 per condition. (f) Immunoblotting of lysates prepared from *Lrp1* control and cKO^OL^ OPC/OL cultures after 3 days in DM. Representative blots were probed with anti-p-AKT (S473), anti-p-tAKT, anti-pERK (1/2), anti-tERK (1/2), and anti-GAPDH. (g) Quantification of protein levels detected by immunoblotting per OPC/OL culture from *Lrp1* control and cKO^OL^ mice. pAKT/AKT, n= 5 for *Lrp1* control and cKO^OL^ mice; pERK/tERK, n= 3 for *Lrp1* control and cKO^OL^ mice. Results are presented as the mean ± SEM, \**p*<0.05, \*\**p*<0.01, and \*\*\**p*<0.001, Student’s *t*-test. For a detailed statistical report, see Figure3-source data1.

### Loss of *Lrp1* attenuates OPC differentiation *in vitro*

To independently assess the role of *Lrp1* in OL differentiation, we isolated OPCs from brains of *Lrp1* control and cKO^OL^ pups (**Figure 3e**). OPCs were kept in PDGF-AA containing growth medium (GM), allowing them to proliferate or switched to differentiation medium (DM) containing triiodothyronine (T3). Staining of cells for the proliferation marker Ki67 did not reveal any change in OPC proliferation in *Lrp1* cKO^OL^ cultures after 1 or 2 days in GM (**Figure 3-figure supplement 1a-1c**). After 3 days in DM, the number of NG2^+^ and CNP^+^ OLs was comparable between *Lrp1* control and cKO^OL^ cultures (**Figure 3f and 3h**). An abundant signal for LRP1β was detected in *Lrp1* control lysate, but LRP1 was not detectable in *Lrp1* cKO^OL^ cell lysate, demonstrating efficient gene deletion in the OL linage (**Figure 3g**). Moreover, a significant reduction in CNP, MAG, and PLP was detected in *Lrp1* cKO^OL^ cell lysates (**Figure 3-figure supplement 1d and 1e**). As LRP1 signaling is known to regulate Erk1/2 and AKT activity(Yoon et al., 2013), immunoblots were probed for pAKT (S473) and pErk1/2. When normalized to total AKT, levels of pAKT are reduced in *Lrp1* cKO^OL^ lysate, while pErk1/2 levels are comparable between *Lrp1* control and cKO^OL^ lysates (**Figure 3-figure supplement 1f and 1g**). Extended culture of *Lrp1* deficient OLs in DM for 5 days is not sufficient to restore myelin protein levels. Compared to *Lrp1* control cultures, mutants show significantly fewer MAG^+^, PLP^+^, and MBP^+^ cells (**Figure 3i and 3k**) and immunoblotting of cell lysates revealed a reduction in total CNP, MAG, PLP and MBP (**Figure 3j and 3l**). Collectively, our studies demonstrate a cell-autonomous function for *Lrp1* in the OL lineage, important for cell differentiation into myelin sheet producing OLs.

**Figure 4:**
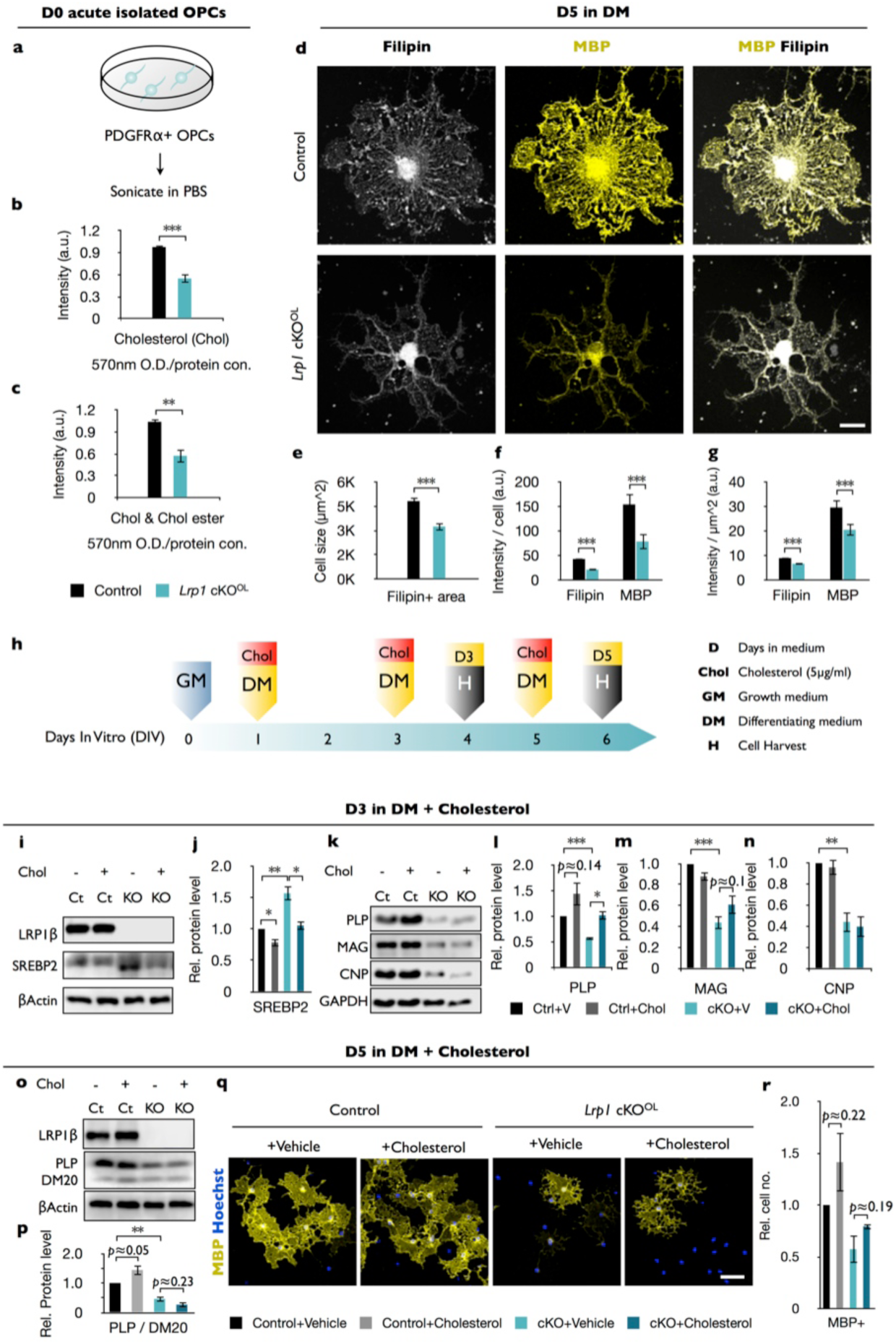
*Lrp1* deficient OPCs show reduced levels of free cholesterol. (a) OPCs were isolated from P10 brains by anti-PDGFRα immunopanning, sonicated and subjected to measurement of free and esterified cholesterol. (b and c) Quantification of free cholesterol (Chol) (b) and total cholesterol (Chol & Chol ester) (c) from *Lrp1* control (n= 5) and cKO^OL^ (n= 5) OPCs. (d) *Lrp1* control and cKO^OL^ OLs after 5 days in DM stained with filipin and anti-MBP. Scale bar= 10μm. (e-g) Quantification of OL size in μm^2^ (e), the intensity of filipin and MBP labeling per cell (f), and the intensity of filipin and MBP staining per μm^2^ (g). For *Lrp1* control and cKO^OL^ OLs, n= 29 cells from 3 mice in each group. (h) Timeline in days showing when growth medium (GM) or differentiation medium (DM) with cholesterol was added and when cells were harvested. (i and k) Immunoblotting of OL lysates prepared from *Lrp1* control and cKO^OL^ cultures after 3 days in DM. Representative blots were probed with anti-LRP1β, anti-SREBP2, anti-β-actin, anti-PLP, anti-MAG, anti-CNP, and anti-GAPDH. (j, l-n) Quantification of SREBP2 (j), PLP (l), MAG (m), and CNP (n) in *Lrp1* control and cKO^OL^ cultures with (+) or without (-) bath applied cholesterol. Number of independent immunoblots: anti-PLP, and MAG, n= 3 per condition; anti-SREBP2 and anti-CNP, n= 4 per condition. (o) Immunoblotting of OL lysates prepared from *Lrp1* control and cKO^OL^ cultures after 5 days in DM with (+) or without (-) bath applied cholesterol. Representative blots were probed with anti-LRP1β, anti-PLP/DM20, and anti-β-actin. (p) Quantification of PLP (n=4 per condition) in *Lrp1* control and cKO^OL^ cultures with (+) or without (-) bath applied cholesterol (q) Immunostaining of OLs after 5 days in DM with (+) or without (-) bath applied cholesterol. Primary OLs stained with anti-MBP and Hoechst dye33342. Scale bar= 100μm. (r) Quantification showing relative number of MBP^+^ cells in *Lrp1* control and cKO^OL^ cultures (n= 3 per condition). Results are shown as mean values ± SEM, *p<0.05, **p<0.01, and ***p<0.001, 2-way ANOVA, post hoc *t*-test. For a detailed statistical report, see Figure4-source data1.

### *Lrp1* deficiency in OPC and OLs causes a reduction in free cholesterol

While LRP1 has been implicated in cholesterol uptake and homeostasis in non-neural cell types (van de Sluis et al., 2017), a role in cholesterol homeostasis in the OL-lineage has not yet been investigated. We find that OPCs deficient for *Lrp1* (*Lrp1*^-/-^) have reduced levels of free cholesterol compared to *Lrp1* control OPCs (**Figure 4a,b**). Levels of cholesteryl-ester are very low in the CNS(Bjorkhem and Meaney, 2004) and near the detection limit in the *Lrp1* control and *Lrp1*^*-/-*^ OPCs (**Figure 4c**). Morphological studies with MBP^+^ OLs revealed a significant reduction in myelin-like membrane sheet expansion in *Lrp1*^-/-^ OLs (**Figure 4d and 4e**), reminiscent of wildtype OLs cultures treated with statins to inhibit HMG-CoA reductase, the rate limiting enzyme in the cholesterol biosynthetic pathway(Maier et al., 2009, Paintlia et al., 2010, Smolders et al., 2010). To assess cholesterol distribution in primary OLs, cultures were stained with filipin. In *Lrp1* control OLs, staining was observed on myelin sheets and was particularly strong near the cell soma. In *Lrp1*^-/-^ OLs, filipin and MBP staining was significantly reduced (**Figure 4f**). Reduced filipin staining is not simply a reflection of smaller cell size, as staining intensity was decreased when normalized to myelin sheet surface area (**Figure 4g**). Thus, independent measurements revealed a dysregulation of cholesterol homeostasis in *Lrp1*^-/-^ OPCs/OLs.

**Figure 4-figure supplement 1:**
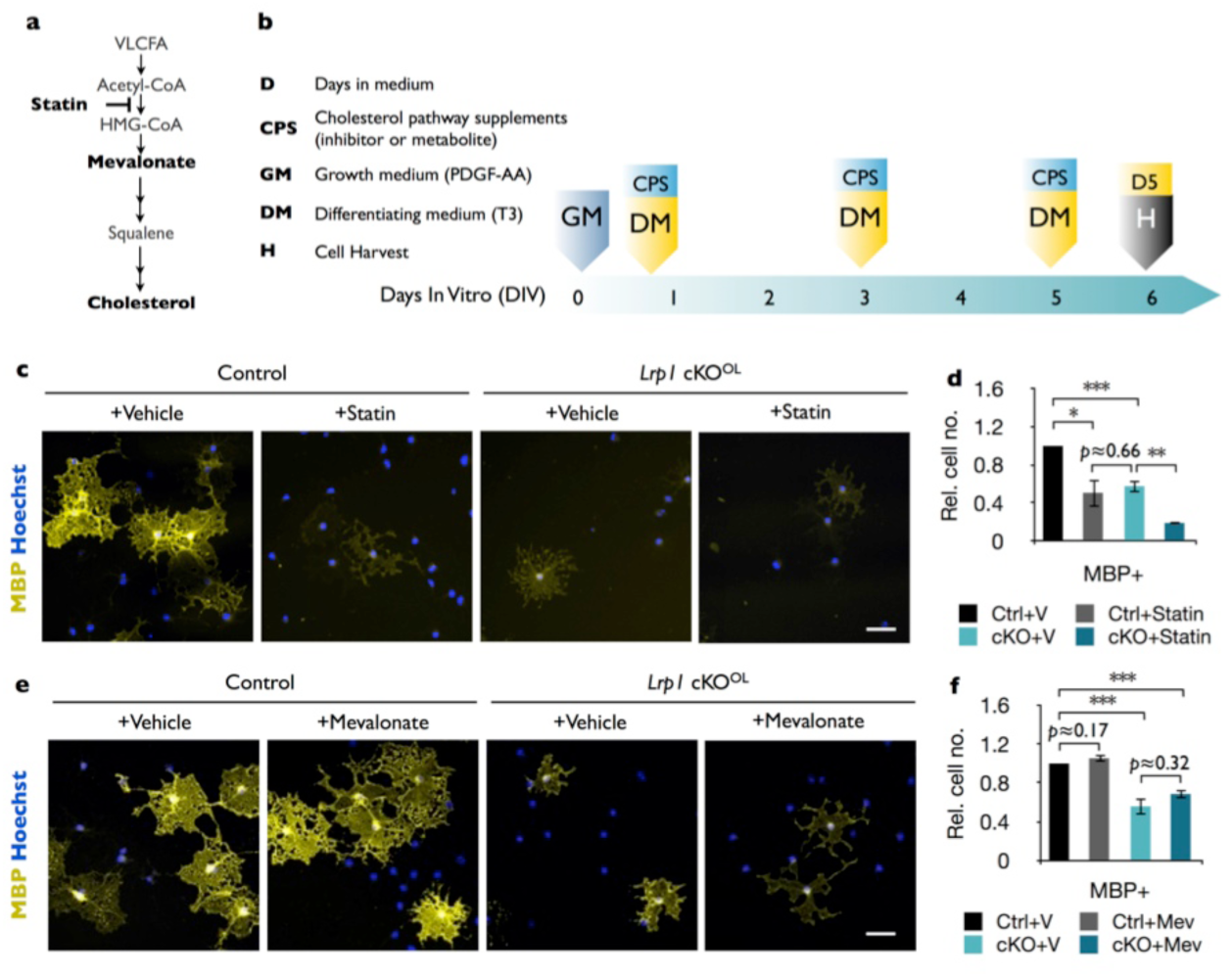
*Lrp1* deficient OLs are sensitive to statin treatment but not to exogenously supplied mevalonate. (a) Cholesterol biosynthetic pathway and site of action of statin (simvastatin). (b) Timeline in days showing when growth medium (GM) and differentiation medium (DM) supplemented with cholesterol pathway supplements (CPS) either simvastatin or mevalonate (Mev) were supplied, and when cells were harvested (H). (c) Immunostaining of OL cultures after 5 days in DM ±statin. Representative cell culture probed with anti-MBP and Hoechst dye33342. Scale bar= 50μm. (d) Quantification of MBP^+^ cells in *Lrp1* control cultures + vehicle (n= 4), *Lrp1* control cultures + statin (n= 3), *Lrp1* cKO^OL^ cultures + vehicle (n= 4), and *Lrp1* cKO^OL^ cultures + statin (n= 3). (e) Immunostaining of OL cultures after 5 days in DM ± mevalonate. Representative cell culture probed with anti-MBP and Hoechst dye33342. Scale bar= 50μm. (f) Quantification of MBP^+^ cells in *Lrp1* control culture + vehicle (n= 3), *Lrp1* control culture + mevalonate (n= 3), *Lrp1* cKO^OL^ culture + vehicle (n= 3), and *Lrp1* cKO^OL^ culture + mevalonate (n= 3). Results are shown as mean values ±SEM, \**p*<0.05, \*\**p*<0.01, and \*\*\**p*<0.001, 2-way ANOVA, post hoc *t*-test. For a detailed statistical report, see Figure4-source data1.

Cellular lipid homeostasis is regulated by a family of membrane-bound basic helix-loop-helix transcription factors, called sterol-regulatory element-binding proteins (SREBPs). Precursors of SREBPs are localized to the ER and activated if cholesterol levels drop below a certain threshold (Goldstein et al., 2006, Faust and Kovacs, 2014). To assess whether *Lrp1* deficiency leads to an increase in SREBP2, OLs were cultured for 3 days in DM and analyzed by immunoblotting. Compared to *Lrp1* control OLs, we observed a strong upregulation of SREBP2 in *Lrp1*^*-/-*^ OLs. Elevated SREBP2 can be reversed to normal levels by exogenous cholesterol directly added to the culture medium (**Figure 4i and 4j**). This shows the existence of LRP1 independent cholesterol uptake mechanisms in *Lrp1*^*-/-*^ OLs and a normal physiological response to elevated levels of cellular cholesterol. In *Lrp1* control cultures, bath application of cholesterol leads to a small, yet significant decrease in SREBP2 (**Figure 4j**). Given the importance of cholesterol in OL maturation(Saher et al., 2005, Kramer-Albers et al., 2006, Mathews et al., 2014), we examined whether the differentiation block of *Lrp1*^*-/-*^ OLs can be rescued by bath-applied cholesterol. Remarkably, treatment of *Lrp1*^*-/-*^ OLs for 3 days with cholesterol failed to elevate PLP, MAG or CNP anywhere near the levels observed in *Lrp1* control OLs (**Figure 4k-n**). Cholesterol treated *Lrp1*^*-/-*^ OLs showed a modest increase in PLP but levels remained below *Lrp1* control OLs. To ask whether prolonged treatment with exogenous cholesterol promotes OL differentiation in *Lrp1*^-/-^ cultures, cells were kept for 5 days in DM, either with or without cholesterol. Similar to the 3 day treatment, the 5 day treatment failed to increase PLP levels (**Figure 4o and 4p**) or the number of MBP^+^ OLs (**Figure 4q and 4r**). While differentiation of *Lrp1*^-/-^ OLs cannot be “rescued” by bath applied cholesterol, cell are highly sensitive to a further reduction in cholesterol, as blocking of cholesterol synthesis with simvastatin leads to a further reduction in MBP^+^ OLs (**Figure 4-figure supplement 1c and 1d**). As cholesterol is only one of many lipid derivatives produced by the cholesterol biosynthetic pathway (**Figure 4-figure supplement 1a**), we tested whether mevalonate, an upstream metabolite in the cholesterol biosynthetic pathway improves OPC differentiation. However, similar to cholesterol, exogenously supplied mevalonate fails to increase differentiation of *Lrp1*^-/-^ OPCs into MBP^+^ OLs (**Figure 4-figure supplement 1e and 1f**). Taken together, *Lrp1* deficiency in the OL-lineage leads to a drop in cellular cholesterol, and bath application of cholesterol or mevalonate is not sufficient to drive differentiation into mature OLs. Our data suggest that in addition to cholesterol homeostasis, LRP1 regulates other biological processes important for OPC differentiation.

**Figure 5-figure supplement 1:**
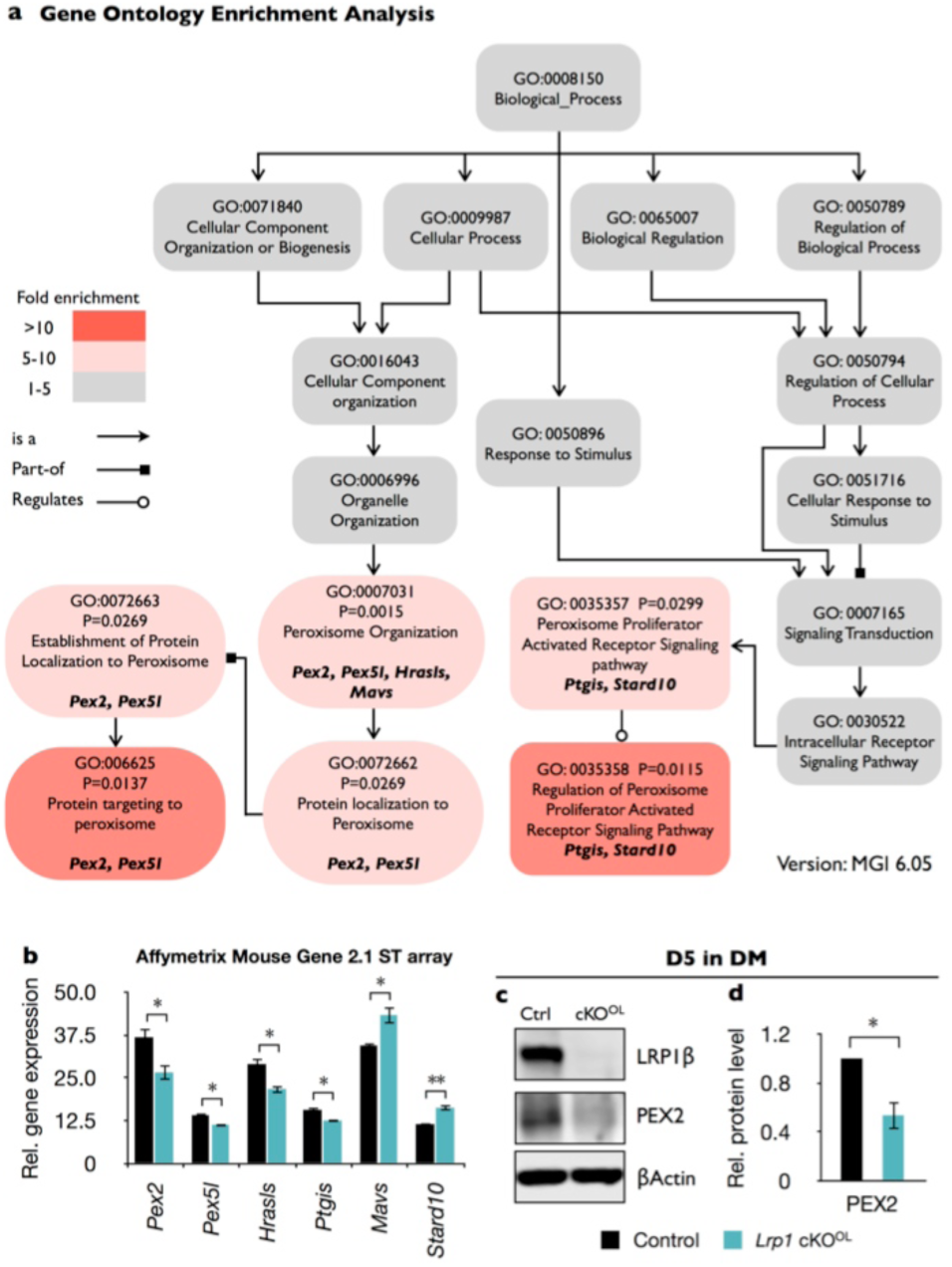
Gene ontology (GO) analysis of *Lrp1* deficient OPCs revealed enrichment of peroxisomal genes. Acutely isolated OPCs from *Lrp1*^*+/+*^ and *Lrp1*^*flox/flox*^;*Olig2-cre* mouse pups were subjected to microarray analysis. (a) GO structure of biological process module related to peroxisome function. Each box shows the GO term ID, *p*-value, GO term, and the genes from the input list associated with the GO term. The color of each box shows the level of enrichment for each GO term. Specific GO terms were queried with the Mouse Genome Informatics (MGI) GO browser. *P*-values were calculated by Fisher’s exact test. The fold enrichment was calculated by dividing the ratio of genes that are associated with each GO term from the input list by the ratio of genes that are expected in the database. (b) Quantification showing relative expression level of gene products that are associated with specific GO terms listed in (a). Gene products were prepared from acutely isolated OPCs for *Lrp1* controls (n= 4) and cKO^OL^ (n =4) and analyze by Affymetrix mouse gene 2.1 ST array. Differentially regulated gene products include *Pex2* (peroxisomal biogenesis factor 2), *Pex5l* (peroxisomal biogenesis factor 5 like), *Hrasls* (hRas-like suppressor), *Ptgis* (prostaglandin I2 synthase), *Mavs* (Mitochondrial antiviral signaling), and *Stard10* (StAR-related lipid transfer protein 10). (c) Immunoblotting of lysates prepared from *Lrp1* control and cKO^OL^ cultures after 5 days in DM. Representative blots were probed with anti-LRP1β, anti-Pex2, and anti-β-actin. (d) Quantification of Pex2 in *Lrp1* control (n= 3) and cKO^OL^ (n= 3) cultures. Results are shown as mean values ±SEM, *p<0.05 and **p<0.01, Student’s *t*-test. For a detailed statistical report, see Figure5-source data1.

### *Lrp1* deficiency impairs peroxisome biogenesis

To further investigate what type of biological processes might be dysregulated by *Lrp1* deficiency, we performed transcriptomic analyses of OPCs acutely isolated from *Lrp1* control and cKO^OL^ pups. Gene ontology (GO) analysis identified differences in “peroxisome organization” and “peroxisome proliferation-associated receptor (PPAR) signaling pathway” (**Figure 5-figure supplement 1a**). Six gene products regulated by *Lrp1* belong to peroxisome and PPAR GO terms, including *Pex2*, *Pex5l*, *Hrasls*, *Ptgis*, *Mavs*, and *Stard10* (**Figure 5-figure supplement 1b**). Western blot analysis of *Lrp1*^-/-^ OLs further revealed a significant reduction in PEX2 after 5 days in DM (**Figure 5-figure supplement 1c and 1d**). Because PEX2 has been implicated in peroxisome biogenesis (Gootjes et al., 2004), and peroxisome biogenesis disorders (PBDs) are typically associated with impaired lipid metabolism and CNS myelin defects(Krause et al., 2006), this prompted us to further explore a potential link between LRP1 and peroxisomes. To assess whether the observed reduction in PEX2 impacts peroxisome density in primary OLs after 5 days in DM, MBP^+^ OLs were stained with anti-PMP70, an ATP-binding cassette transporter enriched in peroxisomes (**Figure 5a**). In *Lrp1*^*-/-*^ OLs, we observed reduced PMP70 staining (**Figure 5b**) and a decrease in the total number of peroxisomes (**Figure 5c**). Normalization of peroxisome counts to cell size revealed that the reduction in *Lrp1*^-/-^ OLs is not simply a reflection of smaller cells (**Figure 5d**).

**Figure 5:**
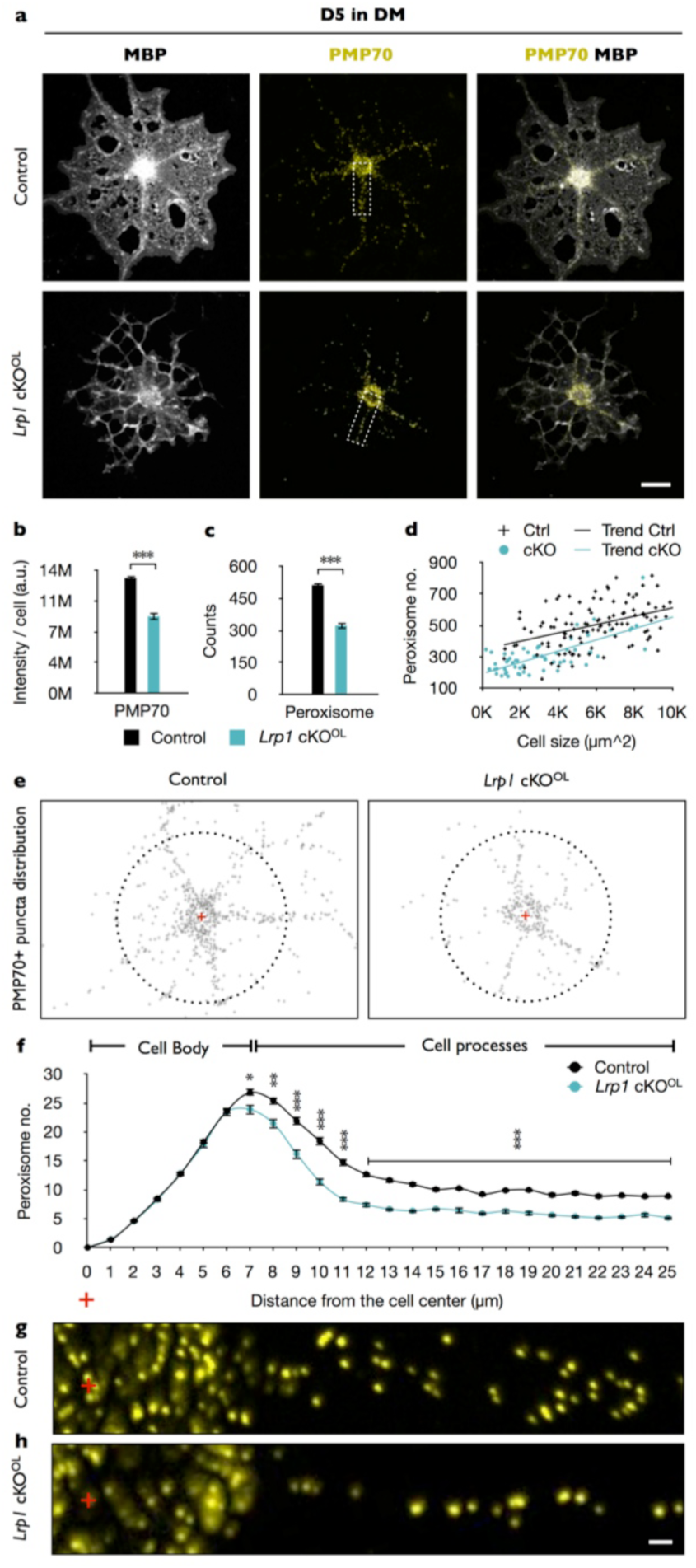
In the OL-linage loss of *Lrp1* leads to a reduction in peroxisomes. (a) Primary OLs prepared from *Lrp1* control and cKO^OL^ OL pups, cultured for 5 days in DM were stained with anti-MBP and anti-PMP70. Scale bar= 10μm. (b-d) Quantification of PMP70 labeling intensity per cell (b), PMP70^+^ puncta per cell (c), and scatter plot showing the number of PMP70^+^ peroxisomes as a function of cell size for MBP^+^ OLs of *Lrp1* control and *Lrp1* cKO^OL^ cultures. (d). For *Lrp1* control OLs, n= 112 cells from 3 mice. For *Lrp1* cKO^OL^ OLs, n= 60 cells from 3 mice. (e) Representative distribution of the PMP70^+^ puncta of *Lrp1* control and cKO^OL^ cells. The center of the cell is marked with a red cross. Puncta within a 25μm radius from the center (dashed circle) were subjected to quantification. (f) Quantification of peroxisome number plotted against the distance from the center of *Lrp1* control (n= 113 cells, 3 mice) and cKO^OL^ (n= 63 cells, 3 mice) OLs. (g and h) Representative high magnification views of PMP70^+^ puncta from white dashed box in (a) aligned with (f). Scale bar= 1μm. Results are shown as mean values ± SEM, *p<0.05, **p<0.01, and ***p<0.001, Student’s *t*-test. For a detailed statistical report, see Figure5-source data1.

The peroxisome is a highly dynamic organelle, comprised of over 50 enzymes, many of which participate in lipid metabolism, including the pre-squalene sequence of the cholesterol biosynthetic pathway (Faust and Kovacs, 2014). The subcellular localization of peroxisomes is thought to be important for ensuring a timely response to metabolic demands (Berger et al., 2016). This prompted us to analyze the subcellular distribution of peroxisomes in primary OLs. Interestingly, while the number of PMP70 positive puncta near the cell soma is comparable between *Lrp1* control and *Lrp1*^-/-^ OLs, we observed a significant drop in peroxisomes along radial processes of MBP^+^ OLs (**Figure 5e-5h**).

**Figure 6:**
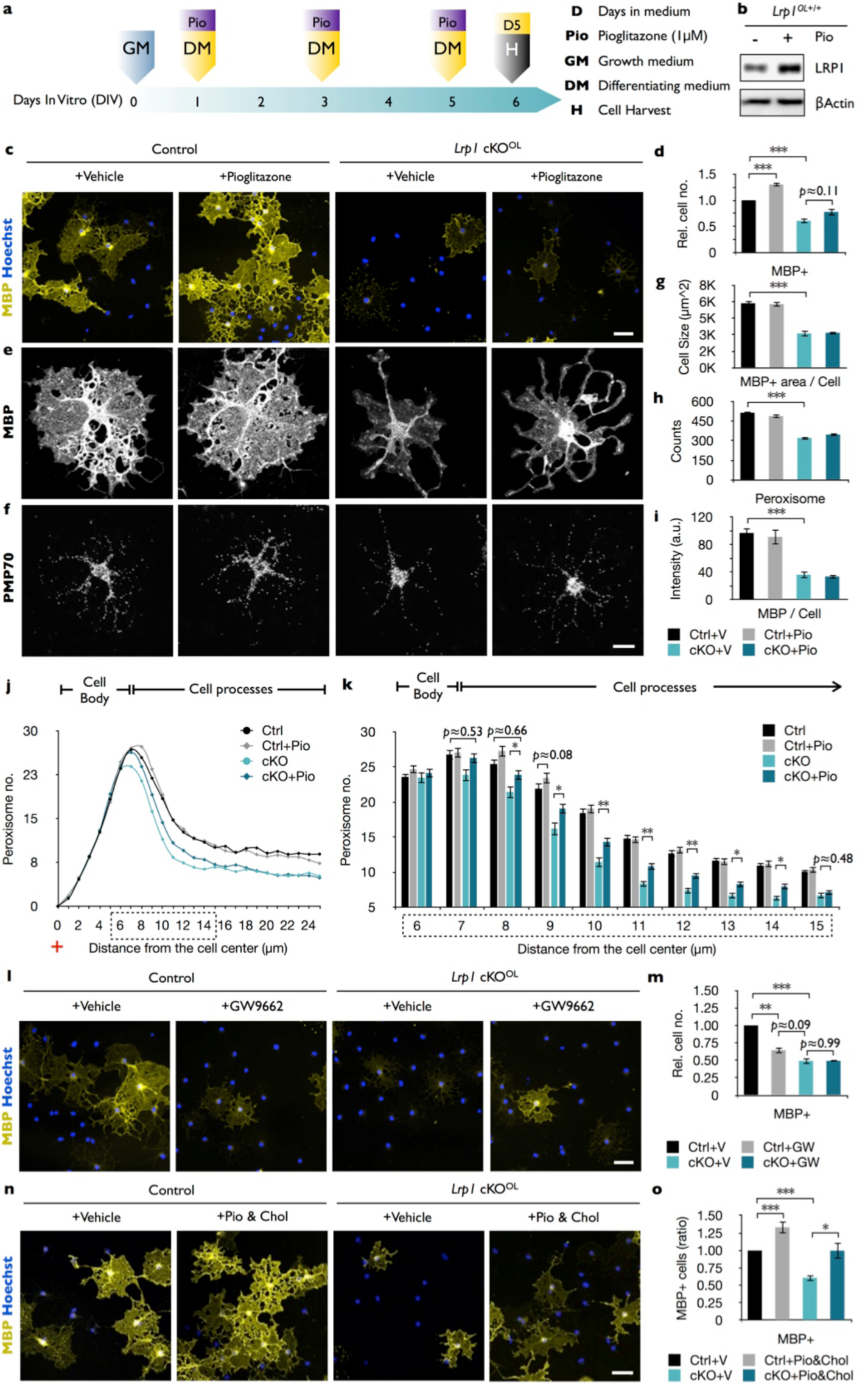
Peroxisome proliferator and cholesterol combined treatment rescues OL differentiation in *Lrp1* deficient OLs. (a) Timeline in days showing when growth medium (GM) or differentiation medium (DM) with pioglitazone (Pio) were supplied and cells were harvested for analysis. (b) Immunoblots of OL lysates prepared from *Lrp1* control and cKO^OL^ cultures after 5 days in DM, probed with anti-LRP1β and anti-β-actin. (c) Immunostaining of *Lrp1* control and cKO^OL^ cultures after 5 days in DM. Representative cell culture probed with anti-MBP and Hoechst. Scale bar= 50μm. (d) Quantification of MBP^+^ cells in *Lrp1* control + vehicle (n= 6), *Lrp1* control + pioglitazone (n= 6), cKO^OL^ + vehicle (n= 4), and cKO^OL^ + pioglitazone (n= 4) treated cultures. (e-f) Primary OLs probed with anti-MBP and anti-PMP70. Scale bar= 10μm. (g-i) Quantification of OL size in μm^2^ (g), the number of PMP70^+^ puncta and (h), the intensity of MBP staining per cell (i). (j) Distribution of peroxisomes as a function of distance from the cell center in *Lrp1* control and cKO^OL^ cultures with (+) or without (-) pioglitazone. The number of PMP70^+^ peroxisomes between 6-15μm in *Lrp1* control and cKO^OL^ cultures was subjected to statistical analysis in (k). *Lrp1* control (n= 112 cells, 3 mice), *Lrp1* control + pio (n= 180 cells, 3 mice), cKO^OL^ (n= 60 cells, 3 mice), and cKO^OL^ + Pio. (n= 110 cells, 3 mice) OLs (k). (l) Immunostaining of OLs after 5 days in DM with (+) or without (-) GW9662, probed with anti-MBP and Hoechst dye33342. Scale bar= 50μm. (m) Quantification of MBP^+^ cells under each of the 4 different conditions (n=3 per condition). (n) Immunostaining of *Lrp1* control and cKO^OL^ cultures after 5 days in DM, probed with anti-MBP and Hoechst dye33342. Scale bar= 50μm. (o) Quantification of MBP^+^ cells in *Lrp1* control + vehicle (n= 4), *Lrp1* control + pioglitazone & cholesterol (n= 3), cKO^OL^ + vehicle (n= 4), and cKO^OL^ + pioglitazone & cholesterol (n= 3) treated cultures. Results are shown as mean values ± SEM, *p<0.05, **p<0.01, and ***p<0.001, 2-way ANOVA, post hoc *t*-test. For a detailed statistical report, see Figure6-source data1.

### Combination treatment of cholesterol and PPARγ agonist rescues the differentiation block in *Lrp1* deficient OPCs

In endothelial cells, the LRP1-ICD functions as a co-activator of PPARγ, a key regulator of lipid and glucose metabolism (Mao et al., 2017). Activated PPARγ moves into the nucleus to control gene expression by binding to PPAR-responsive elements (PPREs) on numerous target genes, including *Lrp1* (Gauthier et al., 2003). In addition, PPREs are found in genes important for lipid and glucose metabolism, and peroxisome biogenesis (Fang et al., 2016, Hofer et al., 2017). *In vitro*, a 5 day treatment of *Lrp1* control OPCs with pioglitazone, an agonist of PPARγ, results in elevated LRP1 (**Figure 6a and 6b**) and accelerated differentiation into MBP^+^ OLs (**Figure 6c and 6d**) (Bernardo et al., 2009). This stands in marked contrast to *Lrp1*^-/-^ cultures, where pioglitazone treatment fails to accelerate OPC differentiation (**Figure 6c and 6d**). Moreover, pioglitazone does not regulate PMP70 staining intensity in *Lrp1* control or *Lrp1*^-/-^ OLs, nor does it have any effect on total peroxisome counts per cell (**Figure 6e-6i**). However, pioglitazone leads to a modest but significant increase in the number of peroxisomes located in cellular processes of *Lrp1*^*-/-*^ OLs (**Figure 6j and 6k**). Treatment of *Lrp1* control OPCs with the PPARγ antagonist GW9662 blocks differentiation into MBP^+^ OLs (Roth et al., 2003), but does not lead to a further reduction in MBP^+^ cells in *Lrp1*^*-/-*^ OL cultures (**Figure 6m**). This suggests that in *Lrp1*^*-/-*^ OLs PPARγ is not active. Given LRP1’s multifunctional receptor role, we asked whether simultaneous treatment with pioglitazone and exogenous cholesterol is sufficient to rescue the differentiation block of *Lrp1*^*-/-*^ OPCs. This is indeed the case, as the number of MBP^+^ cells in *Lrp1*^*-/-*^ cultures is significantly increased by the combination treatment, suggesting an additive effect toward OPC differentiation (**Figure 6n and 6o**). Together, these findings indicate that in OPCs LRP1 regulates multiple metabolic functions important for OL differentiation. In addition to its known role in cholesterol homeostasis, LRP1 regulates expression of PEX2 and thereby metabolic functions associated with peroxisomes.

## Discussion

LRP1 function in the OL-lineage is necessary for proper CNS myelin development and the timely repair of a chemically induced focal white matter lesion *in vivo*. Optic nerves of *Lrp1* cKO^OL^ show fewer myelinated axons, thinning of myelin sheaths, and an increase in nodal defects. Morphological defects have a physiological correlate, as *Lrp1* cKO^OL^ mice exhibit faulty nerve conduction. Mechanistically, *Lrp1* deficiency disrupts multiple signaling pathways implicated in OL differentiation, including AKT activation, cholesterol homeostasis, PPARγ activation, and peroxisome biogenesis. The pleiotropic roles of LRP1 in OPC differentiation are further underscored by the fact that restoring cholesterol homeostasis or activation of PPARγ alone is not sufficient to drive differentiation. Only when cholesterol supplementation is combined with PPARγ activation, differentiation of *Lrp1*^*-/-*^ OPC into MBP^+^ OLs is significantly increased. Taken together, our studies identify a novel link between LRP1 and peroxisomes and suggest that broad metabolic dysregulation in *Lrp1*^*-/-*^ OPCs attenuates differentiation into mature OLs (**Figure 7**).

**Figure 7:**
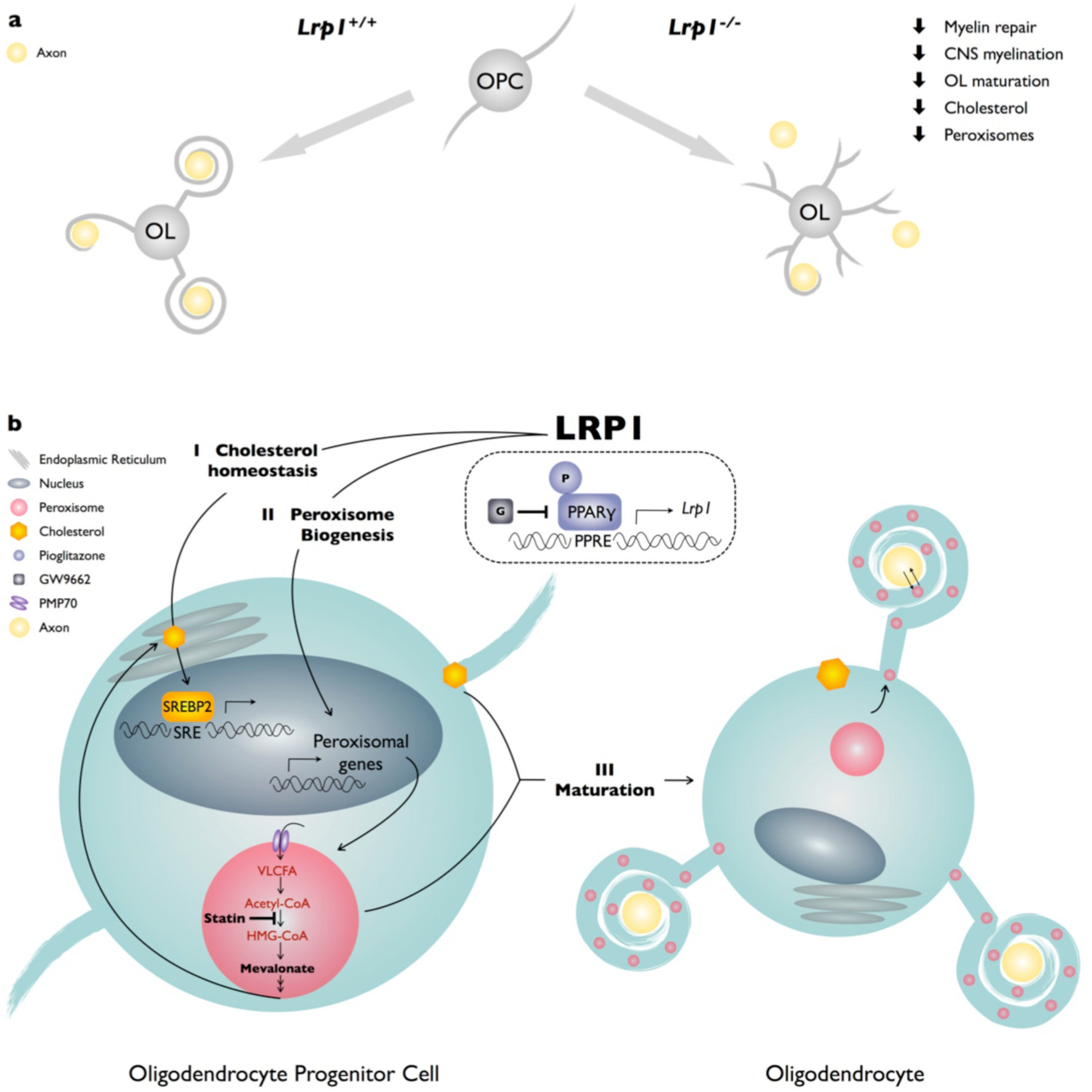
Working model of LRP1 regulated pathways in the OL-lineage. (a) LRP1 function in the OL-lineage is necessary for proper CNS myelin development and the timely repair of a chemically induced focal white matter lesion. In OPCs, *Lrp1* deficiency leads to dysregulation of cholesterol homeostasis and impaired peroxisome biogenesis. (b) LRP1 is a key regulator of multiple pathways important for OPC differentiation into mature myelin producing OLs: I) LRP1 regulates cholesterol homeostatsis; II) LRP1 regulates peroxisome biogenesis; and III) the combined treatment of *Lrp1* deficient primary OPCs with cholesterol and pioglitazone is sufficient to drive maturation into MBP^+^ myelin sheet producing OLs.

In the embryonic neocortex, LRP1 is strongly expressed in the ventricular zone and partially overlaps with nestin^+^ cells, suggesting expression in undifferentiated neural stem and precursor cells (NSPCs) (Hennen et al., 2013). In *Lrp1*^*flox/flox*^ neurospheres, conditional ablation of *Lrp1* reduces cell proliferation and survival, and also negatively impacts differentiation into neurons and O4^+^ OLs, while astrocyte production is significantly increased (Safina et al., 2016). In line with these observations, ablation of *Lrp1* with the *Olig2-cre* driver attenuates OPC differentiation, but neither neurogenesis nor astrocyte production are significantly altered in these mice. A likely explanation for these discrepancies is the OL lineage restricted gene ablation in *Lrp1* cKO^OL^ mice. Studies with highly purified OPCs *in vitro* and OL-linage specific ablation of *Lrp1 in vivo*, both during development and adult white matter repair, suggest a cell-autonomous role for *Lrp1* in OPC maturation and differentiation into myelin producing OLs. Global inducible ablation of *Lrp1* at P5 or at P56 with the *CAG-creER™* driver achieves a ∼40% reduction of LRP1 protein in the brain. Despite multiple daily injections of TM via different routes at P5 or i.p. in adult mice, we were unable to achieve a greater than 40% decrease in LRP1, yet this partial ablation is sufficient to negatively impact myelin development and adult white matter repair. LRP1 is elevated in astrocytes and myeloid cells near multiple sclerosis (MS) lesions (Chuang et al., 2016). Deletion of *Lrp1* in microglia worsens the course of experimental autoimmune encephalomyelitis (EAE) and it has been proposed that loss of *Lrp1* in microglial leads to a proinflammatory phenotype and disease exacerbation (Chuang et al., 2016). Thus, reduced white matter repair in *Lrp1 iKO* mice injected with LPC may be due to loss of *Lrp1* in OPCs, an increase in proinflammatory microglia or a combination thereof. However, studies with *Lrp1 iKO*^*OL*^ mice clearly show that in the adult brain, *Lrp1* function in the OL lineage is necessary for the timely repair of a demyelination lesion. Additional studies, including cell type specific ablation in microglia, astrocytes, and neurons are needed to determine whether *Lrp1* is required in other neural cell types for the repair of LPC inflicted white matter lesions.

Cholesterol does not cross the blood-brain-barrier (Saher and Stumpf, 2015) and CNS resident cells need to either synthesize their own cholesterol or acquire it through horizontal transfer from neighboring cell types, including astrocytes (Camargo et al., 2017). In the OL lineage cholesterol is essential for cell maturation, including myelin gene expression, myelin protein trafficking, and internode formation (Saher et al., 2005, Kramer-Albers et al., 2006, Mathews et al., 2014). Sterol biosynthesis is in part accomplished by peroxisomes. Specifically, the pre-squalene segment of the cholesterol biosynthetic pathway takes place in peroxisomes. However, cholesterol is only one of many lipid derivatives produced by this pathway (Faust and Kovacs, 2014). A drop in intracellular cholesterol leads to an increase in SREBPs, a family of transcription factors that regulate expression of gene products involved in cholesterol and fatty acid synthesis (Goldstein et al., 2006, Faust and Kovacs, 2014). In Schwann cells, SREBPs, and the SREBP activating protein SCAP, are required for AKT/mTOR dependent lipid biosynthesis, myelin membrane synthesis, and normal PNS myelination (Verheijen et al., 2009, Norrmen et al., 2014). In the OL linage blockage of SREBP activation inhibits CNS myelination (Camargo et al., 2017, Monnerie et al., 2017). Blocking of SREBP processing in primary OLs leads to a drop in cellular cholesterol and inhibits cell differentiation and membrane expansion. This can be rescued by addition of cholesterol to the culture medium (Monnerie et al., 2017). In primary OLs, *Lrp1* deficiency leads to activation of SREBP2, yet cells seem to be unable to maintain cholesterol homeostasis, suggesting more global metabolic deficits. The cholesterol sensing apparatus in *Lrp1* deficient OPCs appears to be largely intact, as bath applied cholesterol leads to a reduction in SREBP2. As SREBP2 can be induced by ER stress (Faust and Kovacs, 2014), reversibility of elevated SREBP2 by bath applied cholesterol suggests that *Lrp1* cKO^OL^ cultures upregulate SREBP2 due to cholesterol deficiency and not due to an elevated ER stress response (Faust and Kovacs, 2014). Significantly, restoring cellular cholesterol homeostasis in *Lrp1*^*-/-*^ OPC is not sufficient to overcome the differentiation block, suggesting more widespread functional deficits.

Members of the PPAR subfamily, including PPARα, PPARβ/d, and PPARγ, are ligand-activated transcription factors that belong to the nuclear hormone receptor family (Michalik et al., 2006). PPARs regulate transcription through heterodimerization with the retinoid X receptor (RXR). When activated by a ligand, the dimer modulates transcription via binding to a PPRE motif in the promoter region of target genes (Michalik et al., 2006). A critical role for PPARγ in OL differentiation is supported by the observation that activation with pioglitazone or rosiglitazone accelerates OPC differentiation into mature OLs (Saluja et al., 2001, Roth et al., 2003, Bernardo et al., 2009, De Nuccio et al., 2011, Bernardo et al., 2013) and inhibition with GW9662 blocks OL differentiation. Deficiency for the PPARγ-coactivator-1 alpha (PGC1a) leads to impaired lipid metabolism, including an increase in very long chain fatty acids (VLCFAs) and disruption of cholesterol homeostasis (Xiang et al., 2011, Camacho et al., 2013). In addition, PGC1a deficiency results in defects of peroxisome-related gene function, suggesting the increase in VLCFAs and drop in cholesterol reflects impaired peroxisome function (Baes and Aubourg, 2009). Following γ-secretase dependent processing, the LRP1 ICD can translocate to the nucleus where it associates with transcriptional regulators (May et al., 2002, Carter, 2007). In endothelial cells, the LRP1-ICD binds directly to the nuclear receptor PPARγ to regulate gene products that function in lipid and glucose metabolism (Mao et al., 2017). Treatment of *Lrp1*^*-/-*^ OPCs with pioglitazone leads to an increase in peroxisomes in OL processes but fails to promote differentiation into myelin sheet producing OLs. In the absence of the LRP1-ICD, pioglitazone may fail to fully activate PPARγ (Mao et al., 2017), but the observed increase in PMP70^+^ peroxisomes in OL processes of *Lrp1* deficient cultures suggests that mutant cells can respond to pioglitazone. Because *Lrp1* cKO^OL^ cultures are cholesterol deficient and the LRP1-ICD participates in PPARγ regulated gene expression, we examined whether a combination treatment of cholesterol and pioglitazone rescues the differentiation block in *Lrp1* deficient OPCs/OLs. This was indeed the case, suggesting that *Lrp1* deficiency leads to dysregulation of multiple pathways important for OPC differentiation.

The importance of peroxisomes in the human nervous system is underscored by inherited disorders caused by complete or partial loss of peroxisome function, collectively described as Zellweger spectrum disorders (Berger et al., 2016, Waterham et al., 2016). PEX genes encode peroxins, the proteins required for normal peroxisome assembly and when mutated can cause peroxisome biogenesis disorder (PBD), characterized by a broad range of symptoms, including aberrant brain development, white matter abnormalities, and neurodegeneration (Berger et al., 2016). The genetic basis for PBD is a single mutation in one of the 14 PEX genes, typically leading to deficiencies in numerous metabolic functions carried out by peroxisomes (Steinberg et al., 1993). In developing OLs, *Lrp1* deficiency leads to a decrease in peroxisomal gene products, most prominently a >50% reduction in PEX2, an integral membrane protein that functions in the import of peroxisomal matrix proteins. Mice deficient for *Pex2* lack normal peroxisomes but do assemble empty peroxisome membrane ghosts (Faust and Hatten, 1997). *Pex2* mutant mice show significantly lower plasma cholesterol levels and in the brain the rate of cholesterol synthesis is significantly reduced (Faust and Kovacs, 2014). Recent evidence shows that PBDs not only lead to defects in lipid metabolism, but may also lead to dysregulation of carbohydrate metabolism (Wangler et al., 2017). Mounting evidence points to a close interaction of peroxisomes with other organelles, mitochondria in particular, and disruption of these interactions may underlie the far reaching metabolic defects observed in PBD and genetically manipulated model organisms deficient for a single PEX (Fransen et al., 2017, Wangler et al., 2017). Our studies provide a potential new link between peroxisomes and LRP1 and suggest a new mechanism for how *Lrp1* deficiency may lead to impaired lipid and carbohydrate metabolism.

## Methods

### Mice

All animal handling and surgical procedures were performed in compliance with local and national animal care guidelines and were approved by the Institutional Animal Care and Use Committee (IACUC). *Lrp1*^*flox/flox*^ mice were obtained from Steven Gonias (Stiles et al., 2013) and crossed with *Olig2-Cre* (Schuller et al., 2008), *CAG-CreER™* (#004682, Jackson Laboratories), and *Pdgfrα-CreER™* (Kang et al., 2010) mice. For inducible gene ablation in adult male and female mice, 3 intraperitoneal (i.p.) injections of tamoxifen (75 mg/kg) were given every 24 hrs. Tamoxifen (10 mg/ml) was prepared in a mixture of 9% ethanol and 91% sunflower oil. For inducible gene ablation in juvenile mice, 2 injections in the stomach of 4-hydroxytamoxifen (4OH-TM, 15 μg/g) were given 24 hrs apart. A 10 mg/ml stock solution of 4OH-TM was prepared in 100% ethanol. Mice were kept on a mixed background of C57BL/6J and 129SV. Throughout the study, male and female littermate animals were used. *Lrp1* “control” mice harbor at least one functional *Lrp1* allele. Any of the following genotypes *Lrp1*^*+/+*^, *Lrp1*^*+/flox*^, *Lrp1*^*flox/flox*^, or *Lrp1*^*flox/+*^;*cre*^*+*^ served as *Lrp1* controls

### Genotyping

To obtain genomic DNA (gDNA), tail biopsies were collected, boiled for 30 min in 100μl alkaline lysis buffer (25mM NaOH and 0.2mM EDTA in ddH2O) and neutralized by adding 100μl of 40mM Tris-HCI (pH 5.5). For PCR genotyping, 1-5μl of gDNA was mixed with 0.5μl of 10mM dNTP mix (Promega, C1141), 10μl of 25mM MgCl2, 5μl of 5X Green GoTaq^®^ Buffer (Promega, M791A), 0.2μl of GoTaq^®^ DNA polymerase (Promega, M3005), 0.15μl of each PCR primer stock (90μM), and ddH2O was added to a total volume of 25μl. The following cycling conditions were used: DNA denaturing step (94°C for 3 min) 1X, amplification steps (94°C for 30 sec, 60°C for 1min, and 72°C for 1min) 30X, followed by an elongation step (72°C for 10min) then kept at 4°C for storage. The position of PCR primers used for genotyping is shown in **Figure 1-figure supplement 1**. *Lrp1* WT and loxP-flanked (floxed) alleles were amplified with the forward primer [Lrp1tF10290, F2] 5’-CAT ACC CTC TTC AAA CCC CTT G-3’ and the reverse primer [Lrp1tR10291, R2] 5’-GCA AGC TCT CCT GCT CAG ACC TGG A-3’. The WT allele yields a 291-bp product and the floxed allele yields a 350-bp product. The recombined *Lrp1* allele was amplified with the forward primer [Lrp1rF, F1] 5’-CCC AAG GAA ATC AGG CCT CGG C-3’ and the reverse primer [F2], resulting in a 400-bp product (Hennen et al., 2013). For detection of *Cre*, the forward primer [oIMR1084, CreF] 5’-GCG GTC TGG CAG TAA AAA CTA TC-3’ and reverse primer [oIMR1085, CreR] 5’-GTG AAA CAG CAT TGC TGT CACTT-3’ were used, resulting in a ∼200-bp product. As a positive control, the forward primer [oIMR7338, Il-2pF] 5’-CTA GGC CAC AGA ATT GAA AGA-3’ and the reverse primer [oIMR7339, Il-2pR] 5’-GTA GGT GGA AAT TCT AGC ATC-3’ were mixed with CreF and CreR primers in the same reaction, this reaction yields a 324-bp product (The Jackson laboratory).

### Stereotaxic injection

Male and female mice at postnatal-day (P) 42-56 were used for stereotaxic injection of L-α-Lysophosphatidylcholine (LPC) (Sigma, L4129) into the corpus callosum. Mice were anesthetized with 4% isoflurane mixed with oxygen, mounted on a Stoeling stereotaxic instrument (51730D), and kept under 2% isoflurane anesthesia during surgery. A 5μl-hamilton syringe was loaded with 1% LPC in PBS (Gibco, 10010023), mounted on a motorized stereotaxic pump (Quintessential Stereotaxic injector, 53311) and used for intracranial injection at the following coordinates, AP: 1.25mm, LR: ±1mm, D: 2.25mm. Over a duration of 1 min, 0.5μl of 1% LPC solution was injected on the ipsilateral site and 0.5μl PBS on the contralateral side. After the injection was completed, the needle was kept in place for 2 min before retraction. Following surgery, mice were treated with 3 doses of 70μl of buprenorphine (0.3 mg/ml) every 12 hours. Brains were collected at day 10, and 21 post injection.

### Histochemistry

Animals were deeply anesthetized with a mixture of ketamine/xylazine (25mg/ml ketamine and 2.5mg/ml xylazine in PBS) and perfused trans-cardially with ice-cold PBS for 5 min, followed by ice-cold 4% paraformaldehyde in PBS (4%PFA/PBS) for 5 min. Brains were harvested and post-fixed for 2 hours in perfusion solution. Optic nerves were harvested separately and post-fixed for 20 min in perfusion solution. Brains and optic nerves were cryoprotected overnight in 30% sucrose/PBS at 4°C, embedded in OCT (Tissue-Tek, 4583), and flash frozen on dry ice. Serial sections were cut at 20μm (brain tissue) and 10μm (optic nerves) at -20°C using a Leica CM 3050S Cryostat. Serial sections were mounted onto Superfrost^+^ microscope slides (Fisherbrand, 12-550-15) and stored at -20°C.

### *In Situ* Hybridization

Tissue sections mounted on microscope slides were post-fixed overnight in 4%PFA/PBS at 4°C. Sections were then rinsed 3 times for 5 min each in PBS and the edge of microscope slides was demarcated with a DAKO pen (DAKO, S2002). Sections were subsequently incubated in a series of ethanol/water mixtures: 100% for 1min, 100% for 1min, 95% for 1min, 70% for 1min, and 50% for 1min. Sections were then rinsed in 2x saline-sodium citrate (SSC, 150mM NaCl, and 77.5mM sodium citrate in ddH2O, pH7.2) for 1min, and incubated at 37°C for 30 min in proteinase K solution (10μg/ml proteinase K, 100mM Tris-HCl pH8.0, and 0.5mM EDTA in ddH2O). Proteinase digestion was stopped by rinsing sections in ddH2O and then in PBS for 5min each. To quench RNase activity, slides were incubated in 1% triethanolamine (Sigma, 90278) and 0.4% acetic anhydride (Sigma, 320102) mixture in ddH2O for 10 min at room temperature, rinsed once in PBS for 5min and once in 2X SSC for another 5min. To reduce non-specific binding of cRNA probes, sections were pre-incubated with 125μl hybridization buffer (10% Denhardts solution, 40mg/ml baker’s yeast tRNA, 5mg/ml sheared herring sperm DNA, 5X SSC, and 50% formamide in ddH2O) for at least 2 hours at room temperature. Digoxigenin-labeled cRNA probes were generated by run-off *in vitro* transcription as described (Winters et al., 2011a). Anti-sense and sense cRNA probes were diluted in 125μl pre-hybridization buffer to ∼200ng/ml, denatured for 5 min at 85°C, and rapidly cooled on ice for 2 min. Probes were applied to tissue sections, microscope slides covered with parafilm, and incubated at 55°C overnight in a humidified and sealed container. The next morning slides were rinsed in 5X SSC for 1 min at 55°C, 2X SSC for 5 mins at 55°C, and incubated in 0.2X SSC/50% formamide for 30min at 55°C. Sections were rinsed in 0.2X SSC at room temperature for 5min then rinsed with Buffer1 (100mM Tris-HCl pH7.5, and 1.5M NaCl in ddH2O) for 5 min. A 1% blocking solution was prepared by dissolving 1g blocking powder (Roche, 11096176001) in Buffer1 at 55°C, cooled to room temperature (RT), and applied to slides for 1 hour at RT. Slides were rinsed in Buffer1 for 5 min and 125μl anti-Digoxigenin-AP antibody (Roche, 11093274910, 1:2500) in Buffer1 was applied to each slide for 1.5 hours at RT. Sections were rinsed in Buffer1 for 5 min, then rinsed in Buffer2 (100mM Tris-HCl pH9.5, 100mM NaCl, and 5mM MgCl2 in ddH2O) for 5 min, and incubated in alkaline phosphatase (AP) substrate (Roche, 11681451001, 1:50) in Buffer2. The color reaction was developed for 1-48 h and stopped by rinsing sections in PBS for 10 min. Sections were incubated in Hoechst dye 33342 (Life technology, H3570) for 5 min, air dried, mounted with Fluoromount-G^®^ (SouthernBiotech, 0100-01), and dried overnight before imaging under bright-field. The following cRNA probes were used, *Pdgfr* and *Plp* (DNA templates were kindly provided by Richard Lu (Dai et al., 2014)), *Mag* (Winters et al., 2011a), and *Mbp* (a 650-bp probe based on template provided in the Allen Brain Atlas).

### Quantification of lesion size and myelin repair

Serial sections of the corpus callosum, containing the LPC and PBS injection sites were mounted onto glass coverslips and stained by ISH with digoxigenin-labeled cRNA probes specific for *Mbp, Mag, Plp* and *Pdgfra*. For quantification of the white matter lesion area, the same intensity cutoff was set by Image J threshold for all brain sections and used to measure the size of the lesion. The outer rim of the strongly *Mbp*^*+*^ region (lesion^out^) was traced with the ImageJ freehand drawing tool. The inner rim facing the *Mbp*^-^ region (lesion^in^) was traced as well. For each animal examined, the size of the initial lesion area (lesion^out^) in μm^2^ and remyelinated area (lesion^out^-lesion^in^) in μm^2^ was calculated by averaging the measurement from two sections at the lesion core. The lesion core was defined as the section with the largest lesion area (lesion^out^). To determine remyelination, the ratio of (lesion^out^-lesion^in^)/(lesion^out^) in % was calculated. As an initial lesion depth control, criteria of lesion^out^ area must cover the center of the corpus callosum in each serial section set. If a lesion^out^ area was not located within the corpus callosum, the animal and corresponding brain sections were excluded from the analysis.

### Immunostaining

Tissue sections mounted onto microscope slides were rehydrated in PBS for 5 min, permeabilized in 0.1% TritonX-100, and blocked in PHT (1% horse serum and 0.1% TritonX-100 in PBS) for 1 hour at RT. Primary antibodies were diluted in PHT and applied overnight at 4°C. Sections were rinsed in PBS 3 times for 5 min each and appropriate secondary antibodies were applied (Life technologies, Alexa-fluorophore 405, 488, 555, 594, or 647nm, 1:1000). Slides were rinsed in PBS 3 times for 5 min each and mounted with ProLong^®^ Gold antifade reagent (Life technologies, P36930). For quantification of nodal structures, randomly selected fields of view in each nerve were imaged at 96X magnification with an Olympus IX71 microscope, a maximum projection of 6 Z-stacked images of each region was generated, and the stacked images were used for quantification. As axons run in and out of the plane within longitudinal sections, criteria were set to exclude structures in which Caspr staining was unpaired to reduce “false positive” as nodal defect. The following primary antibodies were used: rabbit anti-Olig2 (Millipore, AB9610, 1:500), rat anti-PDGFRα (BD Pharmingen, 558774, 1:500), rabbit anti-GFAP (Dako, Nr. A 0334, 1:2000), mouse anti-APC (Calbiochem, OP80, Clone CC1, 1:500), rabbit anti-Caspr (1:1000, (Peles et al., 1997)), mouse anti-Na Channel (1:75, (Rasband et al., 1999)). For myelin staining, sections were incubated in Fluoromyelin-Green (Life technologies, F34651 1:200) reagent for 15min.

### Transmission electron microscopy (TEM)

Tissue preparation and image acquisition were carried out as described by Winters et al., 2011. Briefly, mice at P10, P21, and P56 were perfused trans-cardially with ice cold PBS for 1 min, followed by a 10 min perfusion with a mixture of 3% PFA and 2.5% glutaraldehyde in 0.1M Sorensen’s buffer. Brains and optic nerves were dissected and post-fixed in perfusion solution overnight at 4°C. Post-fixed brain tissue and optic nerves were rinsed and transferred to 0.1M Sorensen’s buffer and embedded in resin by the University of Michigan Imaging Laboratory Core. Semi-thin (0.5μm) sections were cut and stained with toluidine blue and imaged by light microscopy. Ultra-thin (75nm) sections were cut and imaged at the ultrastructural level with a Philips CM-100 or a JEOL 100CX electron microscope. For each genotype and age at least 3 animals were processed and analyzed. For each animal over 1000 axons in the optic nerve were measured and quantified by ImageJ. For each optic nerve, 10 images at 13,500x magnification were randomly taken and quantified to calculate the g-ratio and the fraction of myelinated axons. The inner (area^in^) and outer (area^out^) rim of each myelin sheath was traced with the ImageJ freehand drawing tool and the area within was calculated. We then derived axon caliber and fiber caliber (2r) by the following: area^in^= r^2^π. The g-ratios were calculated as such: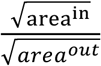. The g-ratio is only accurate if the compact myelin and axon outline can clearly be traced, individual fibers with not clearly define features, i.e. detached myelin were excluded from the quantification.

**Optic nerve CAP recordings** were carried out as described elsewhere(Winters et al., 2011b, Carbajal et al., 2015). Briefly, optic nerves were acutely isolated from P21 mice and transferred into oxygenated ACSF buffer (125 mM NaCl, 1.25mM NaH2PO4, 25mM glucose, 25mM NaHCO3, 2.5mM CaCl2, 1.3mM MgCl2, 2.5mM KCl) for 45 min at RT before transferring into a recording chamber at 37± 0.4°C. Suction pipette electrodes were used for stimulation and recording of the nerve. A computer-driven (Axon pAlamp10.3 software) stimulus isolation unit (WPI, FL) was used to stimulate the optic nerve with 2 mA/50 μs pulses. The recording electrode was connected to a differential AC amplifier (custom-made). A stimulus artifact-subtracting pipette was placed near the recording pipette. A data acquisition system (Axon digidata 1440A, Axon pClamp 10.3, Molecular Devices, CA) was used to digitize the signals. Conduction velocity was calculated from the length of the nerve and the time to peak of each component of the CAP. Amplitudes were normalized to a resistance ratio of 1.7, as described(Fernandes et al., 2014). Raw traces were fitted with 4 Gaussian curves with Origin9.1 software for analysis of individual components of the CAP. Due to limitations in the resolution of individual peaks in short nerves, CAP recordings from nerves that were shorter than 1mm in length were excluded from the analysis.

### OPC/OL primary cultures and drug treatment

OPCs were isolated from P6-P9 mouse pups by anti-PDGFRα (BD Pharmingen, 558774) immunopanning as previously described (Mironova et al., 2016). For plating of cells, 5-7.5 x10^3^ cells (for 12mm cover glass) or 3-5 x10^4^ (12-well plastic plate) were seeded onto PDL pre-coated surface. Primary OPCs were kept in a 10% CO2 incubator at 37°C. To maintain OPCs in a proliferative state, growth medium (20ng/ml PDGF-AA (Peprotech, 100-13A), 4.2μg/ml Forskolin (Sigma, F6886), 10ng/ml CNTF (Peprotech, 450-02), and 1ng/ml NT-3 (Peprotech, 450-03) in SATO) were added to the culture. To induce OPC differentiation, differentiation medium was constituted by adding (4.2μg/ml Forskolin, 10ng/ml CNTF, and 4ng/ml T3 (Sigma, T6397) in SATO) to the culture. For drug treatment, all compounds were mixed with differentiation medium at the desired concentration, and the compound-containing medium was replaced every other day. Stock and working solutions including 20mg/ml cholesterol (Sigma, C8667) in 100% EtOH were kept at RT and warmed up to 37°C before use, then diluted in differentiation medium to 5μg/ml; 10mM pioglitazone (Sigma, E6910) in DMSO was kept at -20°C and diluted in differentiation medium to 1μM; 10mM simvastatin (Sigma, S6196) in DMSO was kept at -20°C and diluted in differentiation medium to 0.5μM; 10mM GW9662 (Sigma, M6191) in DMSO was kept at -20°C and diluted in differentiation medium to 1μM.

### OPC staining and quantification

At different stages of development, OPC/OL cultures were fixed for 15 min in 4%PFA/PBS. Cells were rinsed three times in PBS and permeabilized with 0.1% Triton/PBS solution for 3 mins. Cells were then rinsed in PBS and incubated in blocking solution (3% BSA/PBS) for 1 hour at RT. Primary antibodies were prepared in blocking solution. For immunostaining, 35μl were dropped onto a sheet of parafilm, the coverslips were inverted onto the primary antibody drop, and incubated overnight at 4°C. The following day, coverslips were transferred back to a 24 well-plate and rinsed with PBS 3 times for 5 min each. Secondary antibody ± filipin (Sigma, F9765, 0.1mg/ml) was prepared in blocking solution, 350 μl were added to each well, and the coverslips were incubated for 2 hours at RT. Coverslips then were rinsed in PBS 3 times for 5 min each and stained with Hoechst (1:50,000) for 10s. Coverslips then were rinsed in ddH2O and mounted in ProLong^®^ Gold antifade reagent. For quantification in Figure 3, the % of OL markers^+/^Hoechst^+^ cells was calculated from 10 images that were taken from randomly selected areas in each coverslip at 20X magnification with an Olympus IX71 microscope. For quantification in Figure 4 and after, the % of OL markers+/Hoechst+ cells was calculated from 25 images that were taken from randomly selected areas in each coverslip at 10X magnification with a Zeiss Axio-Observer microscope. For single cell intensity and size measurement in Figures 4-7, individual cell images were taken at 40X magnification with a Zeiss Axio-Observer microscope with Apotom.2. For quantification, the same intensity cutoff was set by Image J threshold to all cells and binary images were generated to define each cell outline. The individual cell outline was applied to original images to measure the intensity of filipin, MBP, or PMP70 staining per cell. For PMP70 puncta distribution analysis, the coordinates of each PMP70+ center were acquired by the process> find maxima function in ImageJ, the cell center coordinate was defined by point selection function, and the distance of each PMP70^+^ dot to the cell center was then calculated. The data were then binned from 1 – 25 mm at 1 mm divisions, and plotted. Primary antibodies included: rat-anti PDGFRα (BD pharmingen, 558774, 1:500), rabbit anti-CNPase (Aves, 27490 R12-2096, 1:500), mouse anti-MAG (Millipore, MAB1567, 1:500), rat anti-MBP (Millipore, MAB386, 1:1000), chicken anti-PLP (Aves, 27592, 1:500), mouse anti-GFAP (Sigma, G3893, 1:1000), chicken anti-GFAP (Aves, GFAP, 1:500), rabbit anti-NG2 (Millipore, AB5320, 1:500), rabbit anti-LRP1β (Abcam, ab92544, 1:500), rabbit anti-PMP70 (Thermo, PA1-650, 1:1000).

### Western blot analysis

Protein lysates were separated by SDS-PAGE and transferred onto PVDF membranes for immunoblotting. Depending on the application, 2 to 10μg of total protein were loaded per well. 2% Blotting-Grade Blocker (Bio-Rad, #170-6404) or 2% BSA fraction V (Fisher, BP1600-100) in 0.1%TBST buffer (0.1% Tween-20, 3M NaCl, 200mM Tris-HCl pH7.4) were used as blocking solutions and membranes were incubated for 1 hour at RT. Primary antibodies were diluted in blocking buffer and used for incubation at 4°C overnight. For protein detection and densitometric analysis, membranes were incubated in Super Signal^®^ West Pico substrate (Thermo, 34080), WesternSure^®^ PREMIUM Chemiluminescent Substrate (LI-COR Biosciences, 926-95000), or Super Signal^®^ West Femto substrate (Thermo, 34095) followed by scanning on a C-DiGit^®^ blot scanner (LI-COR^®^, P/N 3600-00). Images were quantified with Image Studio Lite Western Blot Analysis Software, relative to loading controls. Blots were used for quantification only when the loading control signals were comparable between groups and signals between technical repeats were similar. Primary antibodies included: rabbit anti-LRP1β 85kDa (Abcam, ab92544, 1:2000), mouse anti-βIII tubulin (Promega, G7121, 1:5000), mouse anti-β-actin (Sigma, AC-15 A5441, 1:5000), rat anti-MBP (Millipore, MAB386, 1:1000), rabbit anti-MAG (homemade serum, 1:1000), rabbit anti-PLP (Abcam, ab28486, 1:1000), rat anti-PLP/DM20 (Wendy Macklin AA3 hybridoma, 1:500) rabbit anti-Olig2 (Millipore, AB9610, 1:1000), mouse anti-GFAP (Sigma, G3893, 1:1000), mouse anti-CNPase (Abcam, ab6319, 1:1000), rabbit anti-PXMP3 (PEX2) (One world lab, AP9179c, 1:250), and rabbit anti-SREBP2 (One world lab, 7855, 1:500).

### Cholesterol measurement

OPCs were isolated by immunopanning as described above. OPCs bound to panning plates were collected by scraping with a Scraper (TPP, TP99002) in 250μl of ice-cold PBS and sonicated in an ice-cold water bath (Sonic Dismembrator, Fisher Scientific, Model 500) at 50% amplitude 3 times for 5 sec with a 5 sec interval. The sonicated cell suspensions were immediately used for cholesterol measurement following the manufacturer’s instructions (Chemicon, 428901). For colorimetric detection and quantification of cholesterol, absorbance was measured at 570nm with a Multimode Plate Reader (Molecular Devices, SpectraMax^®^ M5^e^). Results were normalized to total protein concentration measured by DC™ Protein Assay according to the manufacturer’s manual (Bio-Rad, 5000112).

### Microarray and gene ontology analysis

OPCs were isolated by immunopanning as described above and RNA was isolated with the RNeasy Micro Kit (Qiagen, 74004). To compare *Lrp1* control and cKO^OL^ RNA expression profiles, the Mouse Gene ST2.1 Affymetrix array was used. Differentially expressed genes, with a *p*-value <0.05 set as cutoff, were subjected to gene ontology (GO) analysis. Go terms were quarried from Mouse Genome Informatics (MGI) GO browser. The fold enrichment was calculated by dividing the number of genes associated with the GO term in our list by the number of genes associated with the GO term in the database.

### Statistical Analysis

There was no pre-experimental prediction of the difference between control and experimental groups when the study was designed. Therefore, we did not use computational methods to determine sample size a priori. Instead we use the minimum of mice per genotype and experimental treatment for a total of at least 3 independent experiments to achieve the statistical power discussed by Gauch et al (Gauch, 2006). We used littermate *Lrp1* control or *Lrp1* cKO or iKO mice for comparison throughout the study. All independent replicas were biological replicas, rather than technical replicas. For each experiment the sample size (n) is specified in the figure legend. Throughout the study independent replicas (n) indicate biological replica. Technical replicas were used to control for the quality of each measurement and were averaged before quantification and the average value was used as (n= 1) biological replica. Unless indicated otherwise, results are represented as mean value ±SEM. For single pairwise comparison, Student’s *t*-test was used and a *p*-value <0.05 was considered statistically significant. For multiple comparisons, 2-way ANOVA followed by post hoc *t*-test were used. Numbers and R software (see source code file for details) were used for determining statistical significance and graph plotting. For detailed raw data and statistical report, see source data files for each figure. For image processing and quantification, ImageJ 1.47v software was used for threshold setting, annotation, and quantification.

## Acknowledgements

We thank Andy Lieberman for providing *CAG-CreER™* mice, Ben Barres for providing *Olig2-Cre* mice, and Richard Lu for *Plp1* and *Mbp* plasmid DNA. We thank Chang-Ting Lin, Wei-Chin Hou, and Chih-Hsu Lin for bioinformatics consulting. This work was supported by a Bradley Merrill Patten Fellowship (J-PL), the Training Program in Organogenesis T32HD007505 and NIH Cellular and Molecular Biology Training Grant T32-GM007315 (YAM), R01 NS081281 (PS and RJG), the Schmitt Program on Integrative Brain Research (PS), and the Dr. Miriam and Sheldon G. Adelson Medical Foundation on Neural Repair and Rehabilitation (RJG).

## Author contribution

J.-P.L., Y.A.M., P.S., and R.J.G. designed research. J.-P.L., Y.A.M., and R.J.G. performed research. J.-P.L. and P.S. analyzed data. J.-P.L and R.J.G. wrote the paper.

## Additional information

Competing financial interests: the authors declare no competing financial interests.

## References

Ainger K, Avossa D, Morgan F, Hill SJ, Barry C, Barbarese E, Carson JH (1993) Transport and localization of exogenous myelin basic protein mRNA microinjected into oligodendrocytes. J Cell Biol 123:431–441.

Auderset L, Cullen CL, Young KM (2016) Low Density Lipoprotein-Receptor Related Protein 1 Is Differentially Expressed by Neuronal and Glial Populations in the Developing and Mature Mouse Central Nervous System. PLoS One 11:e0155878.

Baes M, Aubourg P (2009) Peroxisomes, myelination, and axonal integrity in the CNS. Neuroscientist 15:367–379.

Berger J, Dorninger F, Forss-Petter S, Kunze M (2016) Peroxisomes in brain development and function. Biochim Biophys Acta 1863:934–955.

Bernardo A, Bianchi D, Magnaghi V, Minghetti L (2009) Peroxisome proliferator-activated receptor-gamma agonists promote differentiation and antioxidant defenses of oligodendrocyte progenitor cells. J Neuropathol Exp Neurol 68:797–808.

Bernardo A, De Simone R, De Nuccio C, Visentin S, Minghetti L (2013) The nuclear receptor peroxisome proliferator-activated receptor-gamma promotes oligodendrocyte differentiation through mechanisms involving mitochondria and oscillatory Ca2+ waves. Biol Chem 394:1607–1614.

Bjorkhem I, Meaney S (2004) Brain cholesterol: long secret life behind a barrier. Arterioscler Thromb Vasc Biol 24:806–815.

Boucher P, Herz J (2011) Signaling through LRP1: Protection from atherosclerosis and beyond. Biochem Pharmacol 81:1–5.

Camacho A, Huang JK, Delint-Ramirez I, Yew Tan C, Fuller M, Lelliott CJ, Vidal-Puig A, Franklin RJ (2013) Peroxisome proliferator-activated receptor gamma-coactivator-1 alpha coordinates sphingolipid metabolism, lipid raft composition and myelin protein synthesis. Eur J Neurosci 38:2672–2683.

Camargo N, Goudriaan A, van Deijk AF, Otte WM, Brouwers JF, Lodder H, Gutmann DH, Nave KA, Dijkhuizen RM, Mansvelder HD, Chrast R, Smit AB, Verheijen MHG (2017) Oligodendroglial myelination requires astrocyte-derived lipids. PLoS Biol 15:e1002605.

Campana WM, Li X, Dragojlovic N, Janes J, Gaultier A, Gonias SL (2006) The low-density lipoprotein receptor-related protein is a pro-survival receptor in Schwann cells: possible implications in peripheral nerve injury. J Neurosci 26:11197–11207.

Carbajal KS, Mironova Y, Ulrich-Lewis JT, Kulkarni D, Grifka-Walk HM, Huber AK, Shrager P, Giger RJ, Segal BM (2015) Th Cell Diversity in Experimental Autoimmune Encephalomyelitis and Multiple Sclerosis. Journal of immunology 195:2552–2559.

Carter CJ (2007) Convergence of genes implicated in Alzheimer’s disease on the cerebral cholesterol shuttle: APP, cholesterol, lipoproteins, and atherosclerosis. Neurochem Int 50:12–38.

Chuang TY, Guo Y, Seki SM, Rosen AM, Johanson DM, Mandell JW, Lucchinetti CF, Gaultier A (2016) LRP1 expression in microglia is protective during CNS autoimmunity. Acta Neuropathol Commun 4:68.

Dai ZM, Sun S, Wang C, Huang H, Hu X, Zhang Z, Lu QR, Qiu M (2014) Stage-specific regulation of oligodendrocyte development by Wnt/beta-catenin signaling. J Neurosci 34:8467–8473.

De Nuccio C, Bernardo A, De Simone R, Mancuso E, Magnaghi V, Visentin S, Minghetti L (2011) Peroxisome proliferator-activated receptor gamma agonists accelerate oligodendrocyte maturation and influence mitochondrial functions and oscillatory Ca(2+) waves. J Neuropathol Exp Neurol 70:900–912.

Emery B, Agalliu D, Cahoy JD, Watkins TA, Dugas JC, Mulinyawe SB, Ibrahim A, Ligon KL, Rowitch DH, Barres BA (2009) Myelin gene regulatory factor is a critical transcriptional regulator required for CNS myelination. Cell 138:172–185.

Fancy SP, Chan JR, Baranzini SE, Franklin RJ, Rowitch DH (2011) Myelin regeneration: a recapitulation of development? Annu Rev Neurosci 34:21–43.

Fang L, Zhang M, Li Y, Liu Y, Cui Q, Wang N (2016) PPARgene: A Database of Experimentally Verified and Computationally Predicted PPAR Target Genes. PPAR Res 2016:6042162.

Faust PL, Hatten ME (1997) Targeted deletion of the PEX2 peroxisome assembly gene in mice provides a model for Zellweger syndrome, a human neuronal migration disorder. J Cell Biol 139:1293–1305.

Faust PL, Kovacs WJ (2014) Cholesterol biosynthesis and ER stress in peroxisome deficiency. Biochimie 98:75–85.

Fernandes KA, Harder JM, John SW, Shrager P, Libby RT (2014) DLK-dependent signaling is important for somal but not axonal degeneration of retinal ganglion cells following axonal injury. Neurobiol Dis 69:108–116.

Fernandez-Castaneda A, Arandjelovic S, Stiles TL, Schlobach RK, Mowen KA, Gonias SL, Gaultier A (2013) Identification of the low density lipoprotein (LDL) receptor-related protein-1 interactome in central nervous system myelin suggests a role in the clearance of necrotic cell debris. J Biol Chem 288:4538–4548.

Franklin RJ, Ffrench-Constant C (2008) Remyelination in the CNS: from biology to therapy. Nat Rev Neurosci 9:839–855.

Fransen M, Lismont C, Walton P (2017) The Peroxisome-Mitochondria Connection: How and Why? Int J Mol Sci 18.

Fuentealba RA, Liu Q, Kanekiyo T, Zhang J, Bu G (2009) Low density lipoprotein receptor-related protein 1 promotes anti-apoptotic signaling in neurons by activating Akt survival pathway. J Biol Chem 284:34045–34053.

Gan M, Jiang P, McLean P, Kanekiyo T, Bu G (2014) Low-density lipoprotein receptor-related protein 1 (LRP1) regulates the stability and function of GluA1 alpha-amino-3-hydroxy-5-methyl-4-isoxazole propionic acid (AMPA) receptor in neurons. PLoS One 9:e113237.

Gauch HG (2006) Winning the Accuracy Game: Three statistical strategies—replicating, blocking and modeling—can help scientists improve accuracy and accelerate progress. American Scientist 94:133–141.

Gauthier A, Vassiliou G, Benoist F, McPherson R (2003) Adipocyte low density lipoprotein receptor-related protein gene expression and function is regulated by peroxisome proliferator-activated receptor gamma. J Biol Chem 278:11945–11953.

Goldstein JL, DeBose-Boyd RA, Brown MS (2006) Protein sensors for membrane sterols. Cell 124:35–46.

Gootjes J, Elpeleg O, Eyskens F, Mandel H, Mitanchez D, Shimozawa N, Suzuki Y, Waterham HR, Wanders RJ (2004) Novel mutations in the PEX2 gene of four unrelated patients with a peroxisome biogenesis disorder. Pediatr Res 55:431–436.

Hennen E, Safina D, Haussmann U, Worsdorfer P, Edenhofer F, Poetsch A, Faissner A (2013) A LewisX glycoprotein screen identifies the low density lipoprotein receptor-related protein 1 (LRP1) as a modulator of oligodendrogenesis in mice. J Biol Chem 288:16538–16545.

Hernandez M, Casaccia P (2015) Interplay between transcriptional control and chromatin regulation in the oligodendrocyte lineage. Glia 63:1357–1375.

Herz J, Clouthier DE, Hammer RE (1992) LDL receptor-related protein internalizes and degrades uPA-PAI-1 complexes and is essential for embryo implantation. Cell 71:411–421.

Hofer DC, Pessentheiner AR, Pelzmann HJ, Schlager S, Madreiter-Sokolowski CT, Kolb D, Eichmann TO, Rechberger G, Bilban M, Graier WF, Kratky D, Bogner-Strauss JG (2017) Critical role of the peroxisomal protein PEX16 in white adipocyte development and lipid homeostasis. Biochim Biophys Acta 1862:358–368.

Kanekiyo T, Bu G (2014) The low-density lipoprotein receptor-related protein 1 and amyloid-beta clearance in Alzheimer’s disease. Front Aging Neurosci 6:93.

Kang SH, Fukaya M, Yang JK, Rothstein JD, Bergles DE (2010) NG2+ CNS glial progenitors remain committed to the oligodendrocyte lineage in postnatal life and following neurodegeneration. Neuron 68:668–681.

Kim J, Yoon H, Basak J, Kim J (2014) Apolipoprotein E in synaptic plasticity and Alzheimer’s disease: potential cellular and molecular mechanisms. Mol Cells 37:767–776.

Kramer-Albers EM, Gehrig-Burger K, Thiele C, Trotter J, Nave KA (2006) Perturbed interactions of mutant proteolipid protein/DM20 with cholesterol and lipid rafts in oligodendroglia: implications for dysmyelination in spastic paraplegia. J Neurosci 26:11743–11752.

Krause C, Rosewich H, Thanos M, Gartner J (2006) Identification of novel mutations in PEX2, PEX6, PEX10, PEX12, and PEX13 in Zellweger spectrum patients. Hum Mutat 27:1157.

Landowski LM, Pavez M, Brown LS, Gasperini R, Taylor BV, West AK, Foa L (2016) Low-density Lipoprotein Receptor-related Proteins in a Novel Mechanism of Axon Guidance and Peripheral Nerve Regeneration. J Biol Chem 291:1092–1102.

Li JS, Yao ZX (2012) MicroRNA patents in demyelinating diseases: a new diagnostic and therapeutic perspective. Recent Pat DNA Gene Seq 6:47–55.

Lillis AP, Van Duyn LB, Murphy-Ullrich JE, Strickland DK (2008) LDL receptor-related protein 1: unique tissue-specific functions revealed by selective gene knockout studies. Physiol Rev 88:887–918.

Liu Q, Trotter J, Zhang J, Peters MM, Cheng H, Bao J, Han X, Weeber EJ, Bu G (2010) Neuronal LRP1 knockout in adult mice leads to impaired brain lipid metabolism and progressive, age-dependent synapse loss and neurodegeneration. J Neurosci 30:17068–17078.

Maier O, De Jonge J, Nomden A, Hoekstra D, Baron W (2009) Lovastatin induces the formation of abnormal myelin-like membrane sheets in primary oligodendrocytes. Glia 57:402–413.

Mantuano E, Jo M, Gonias SL, Campana WM (2010) Low density lipoprotein receptor-related protein (LRP1) regulates Rac1 and RhoA reciprocally to control Schwann cell adhesion and migration. J Biol Chem 285:14259–14266.

Mao H, Lockyer P, Li L, Ballantyne CM, Patterson C, Xie L, Pi X (2017) Endothelial LRP1 regulates metabolic responses by acting as a co-activator of PPARgamma. Nat Commun 8:14960.

Martin AM, Kuhlmann C, Trossbach S, Jaeger S, Waldron E, Roebroek A, Luhmann HJ, Laatsch A, Weggen S, Lessmann V, Pietrzik CU (2008) The functional role of the second NPXY motif of the LRP1 beta-chain in tissue-type plasminogen activator-mediated activation of N-methyl-D-aspartate receptors. J Biol Chem 283:12004–12013.

Mathews ES, Mawdsley DJ, Walker M, Hines JH, Pozzoli M, Appel B (2014) Mutation of 3-hydroxy-3-methylglutaryl CoA synthase I reveals requirements for isoprenoid and cholesterol synthesis in oligodendrocyte migration arrest, axon wrapping, and myelin gene expression. J Neurosci 34:3402–3412.

May P, Bock HH, Herz J (2003) Integration of endocytosis and signal transduction by lipoprotein receptors. Sci STKE 2003:PE12.

May P, Reddy YK, Herz J (2002) Proteolytic processing of low density lipoprotein receptor-related protein mediates regulated release of its intracellular domain. J Biol Chem 277:18736– 18743.

Michalik L, Auwerx J, Berger JP, Chatterjee VK, Glass CK, Gonzalez FJ, Grimaldi PA, Kadowaki T, Lazar MA, O’Rahilly S, Palmer CN, Plutzky J, Reddy JK, Spiegelman BM, Staels B, Wahli W (2006) International Union of Pharmacology. LXI. Peroxisome proliferator-activated receptors. Pharmacol Rev 58:726–741.

Mironova YA, Lenk GM, Lin JP, Lee SJ, Twiss JL, Vaccari I, Bolino A, Havton LA, Min SH, Abrams CS, Shrager P, Meisler MH, Giger RJ (2016) PI(3,5)P biosynthesis regulates oligodendrocyte differentiation by intrinsic and extrinsic mechanisms. Elife 5.

Monnerie H, Romer M, Jensen BK, Millar JS, Jordan-Sciutto KL, Kim SF, Grinspan JB (2017) Reduced sterol regulatory element-binding protein (SREBP) processing through site-1 protease (S1P) inhibition alters oligodendrocyte differentiation in vitro. J Neurochem 140:53–67.

Muratoglu SC, Mikhailenko I, Newton C, Migliorini M, Strickland DK (2010) Low density lipoprotein receptor-related protein 1 (LRP1) forms a signaling complex with platelet-derived growth factor receptor-beta in endosomes and regulates activation of the MAPK pathway. J Biol Chem 285:14308–14317.

Nakajima C, Kulik A, Frotscher M, Herz J, Schafer M, Bock HH, May P (2013) Low density lipoprotein receptor-related protein 1 (LRP1) modulates N-methyl-D-aspartate (NMDA) receptor-dependent intracellular signaling and NMDA-induced regulation of postsynaptic protein complexes. J Biol Chem 288:21909–21923.

Norrmen C, Figlia G, Lebrun-Julien F, Pereira JA, Trotzmuller M, Kofeler HC, Rantanen V, Wessig C, van Deijk AL, Smit AB, Verheijen MH, Ruegg MA, Hall MN, Suter U (2014) mTORC1 controls PNS myelination along the mTORC1-RXRgamma-SREBP-lipid biosynthesis axis in Schwann cells. Cell Rep 9:646–660.

Orita S, Henry K, Mantuano E, Yamauchi K, De Corato A, Ishikawa T, Feltri ML, Wrabetz L, Gaultier A, Pollack M, Ellisman M, Takahashi K, Gonias SL, Campana WM (2013) Schwann cell LRP1 regulates remak bundle ultrastructure and axonal interactions to prevent neuropathic pain. J Neurosci 33:5590–5602.

Paintlia AS, Paintlia MK, Singh AK, Orak JK, Singh I (2010) Activation of PPAR-gamma and PTEN cascade participates in lovastatin-mediated accelerated differentiation of oligodendrocyte progenitor cells. Glia 58:1669–1685.

Peles E, Nativ M, Lustig M, Grumet M, Schilling J, Martinez R, Plowman GD, Schlessinger J (1997) Identification of a novel contactin-associated transmembrane receptor with multiple domains implicated in protein-protein interactions. EMBO J 16:978–988.

Rasband MN, Peles E, Trimmer JS, Levinson SR, Lux SE, Shrager P (1999) Dependence of nodal sodium channel clustering on paranodal axoglial contact in the developing CNS. J Neurosci 19:7516–7528.

Roth AD, Leisewitz AV, Jung JE, Cassina P, Barbeito L, Inestrosa NC, Bronfman M (2003) PPAR gamma activators induce growth arrest and process extension in B12 oligodendrocyte-like cells and terminal differentiation of cultured oligodendrocytes. J Neurosci Res 72:425–435.

Rowitch DH, Kriegstein AR (2010) Developmental genetics of vertebrate glial-cell specification. Nature 468:214–222.

Safina D, Schlitt F, Romeo R, Pflanzner T, Pietrzik CU, Narayanaswami V, Edenhofer F, Faissner A (2016) Low-density lipoprotein receptor-related protein 1 is a novel modulator of radial glia stem cell proliferation, survival, and differentiation. Glia 64:1363–1380.

Saher G, Brugger B, Lappe-Siefke C, Mobius W, Tozawa R, Wehr MC, Wieland F, Ishibashi S, Nave KA (2005) High cholesterol level is essential for myelin membrane growth. Nat Neurosci 8:468–475.

Saher G, Stumpf SK (2015) Cholesterol in myelin biogenesis and hypomyelinating disorders. Biochim Biophys Acta 1851:1083–1094.

Saluja I, Granneman JG, Skoff RP (2001) PPAR delta agonists stimulate oligodendrocyte differentiation in tissue culture. Glia 33:191–204.

Schuller U, Heine VM, Mao J, Kho AT, Dillon AK, Han YG, Huillard E, Sun T, Ligon AH, Qian Y, Ma Q, Alvarez-Buylla A, McMahon AP, Rowitch DH, Ligon KL (2008) Acquisition of granule neuron precursor identity is a critical determinant of progenitor cell competence to form Shh-induced medulloblastoma. Cancer Cell 14:123–134.

Simons M, Lyons DA (2013) Axonal selection and myelin sheath generation in the central nervous system. Curr Opin Cell Biol 25:512–519.

Smolders I, Smets I, Maier O, vandeVen M, Steels P, Ameloot M (2010) Simvastatin interferes with process outgrowth and branching of oligodendrocytes. J Neurosci Res 88:3361–3375.

Steinberg SJ, Raymond GV, Braverman NE, Moser AB (1993) Peroxisome Biogenesis Disorders, Zellweger Syndrome Spectrum. In: GeneReviews(R) (Pagon, R. A. et al., eds) Seattle (WA).

Stiles TL, Dickendesher TL, Gaultier A, Fernandez-Castaneda A, Mantuano E, Giger RJ, Gonias SL (2013) LDL receptor-related protein-1 is a sialic-acid-independent receptor for myelin-associated glycoprotein that functions in neurite outgrowth inhibition by MAG and CNS myelin. J Cell Sci 126:209–220.

Tao W, Moore R, Smith ER, Xu XX (2016) Endocytosis and Physiology: Insights from Disabled-2 Deficient Mice. Front Cell Dev Biol 4:129.

van de Sluis B, Wijers M, Herz J (2017) News on the molecular regulation and function of hepatic low-density lipoprotein receptor and LDLR-related protein 1. Curr Opin Lipidol 28:241– 247.

Verheijen MH, Camargo N, Verdier V, Nadra K, de Preux Charles AS, Medard JJ, Luoma A, Crowther M, Inouye H, Shimano H, Chen S, Brouwers JF, Helms JB, Feltri ML, Wrabetz L, Kirschner D, Chrast R, Smit AB (2009) SCAP is required for timely and proper myelin membrane synthesis. Proc Natl Acad Sci U S A 106:21383–21388.

Wangler MF, Chao YH, Bayat V, Giagtzoglou N, Shinde AB, Putluri N, Coarfa C, Donti T, Graham BH, Faust JE, McNew JA, Moser A, Sardiello M, Baes M, Bellen HJ (2017) Peroxisomal biogenesis is genetically and biochemically linked to carbohydrate metabolism in Drosophila and mouse. PLoS Genet 13:e1006825.

Waterham HR, Ferdinandusse S, Wanders RJ (2016) Human disorders of peroxisome metabolism and biogenesis. Biochim Biophys Acta 1863:922–933.

Winters JJ, Ferguson CJ, Lenk GM, Giger-Mateeva VI, Shrager P, Meisler MH, Giger RJ (2011a) Congenital CNS hypomyelination in the Fig4 null mouse is rescued by neuronal expression of the PI(3,5)P(2) phosphatase Fig4. J Neurosci 31:17736–17751.

Winters JJJ, Ferguson CJC, Lenk GMG, Giger-Mateeva VIV, Shrager PP, Meisler MHM, Giger RJR (2011b) Congenital CNS hypomyelination in the Fig4 null mouse is rescued by neuronal expression of the PI(3,5)P(2) phosphatase Fig4. In: J Neurosci, vol. 31, pp 17736–17751.

Xiang Z, Valenza M, Cui L, Leoni V, Jeong HK, Brilli E, Zhang J, Peng Q, Duan W, Reeves SA, Cattaneo E, Krainc D (2011) Peroxisome-proliferator-activated receptor gamma coactivator 1 alpha contributes to dysmyelination in experimental models of Huntington’s disease. J Neurosci 31:9544–9553.

Yoon C, Van Niekerk EA, Henry K, Ishikawa T, Orita S, Tuszynski MH, Campana WM (2013) Low-density lipoprotein receptor-related protein 1 (LRP1)-dependent cell signaling promotes axonal regeneration. J Biol Chem 288:26557–26568.

Zhang Y, Chen K, Sloan SA, Bennett ML, Scholze AR, O’Keeffe S, Phatnani HP, Guarnieri P, Caneda C, Ruderisch N, Deng S, Liddelow SA, Zhang C, Daneman R, Maniatis T, Barres BA, Wu JQ (2014) An RNA-sequencing transcriptome and splicing database of glia, neurons, and vascular cells of the cerebral cortex. J Neurosci 34:11929–11947.

Zlokovic BV, Deane R, Sagare AP, Bell RD, Winkler EA (2010) Low-density lipoprotein receptor-related protein-1: a serial clearance homeostatic mechanism controlling Alzheimer’s amyloid beta-peptide elimination from the brain. J Neurochem 115:1077–1089.

Zuchero JB, Barres BA (2013) Intrinsic and extrinsic control of oligodendrocyte development. Curr Opin Neurobiol 23:914–920.

